# A metabolic-dysfunction associated steatotic liver acinus biomimetic induces pancreatic islet dysfunction in a coupled microphysiology system

**DOI:** 10.1101/2024.04.22.590598

**Authors:** Julio Aleman, K Ravikumar, Connor Wiegand, Mark E. Schurdak, Lawrence Vernetti, Dillon Gavlock, Celeste Reese, Richard DeBiasio, Greg LaRocca, Yulder Daniel Angarita, Albert Gough, Alejandro Soto-Gutierrez, Jaideep Behari, Vijay Yechoor, Mark T. Miedel, Andrew M. Stern, Ipsita Banerjee, D. Lansing Taylor

## Abstract

Preclinical and clinical studies suggest that lipid-induced hepatic insulin resistance is a primary defect that predisposes to dysfunction in pancreatic islets, implicating a perturbed liver-pancreas axis underlying the comorbidity of T2DM and MASLD. To investigate this hypothesis, we developed a human biomimetic microphysiological system (MPS) coupling our vascularized liver acinus MPS (vLAMPS) with primary islets on a chip (PANIS) enabling MASLD progression and islet dysfunction to be quantitatively assessed. The modular design of this system (vLAMPS-PANIS) allows intra-organ and inter-organ dysregulation to be deconvoluted. When compared to normal fasting (NF) conditions, under early metabolic syndrome (EMS) conditions, the standalone vLAMPS exhibited characteristics of early stage MASLD, while no significant differences were observed in the standalone PANIS. In contrast, with EMS, the coupled vLAMPS-PANIS exhibited a perturbed islet-specific secretome and a significantly dysregulated glucose stimulated insulin secretion (GSIS) response implicating direct signaling from the dysregulated liver acinus to the islets. Correlations between several pairs of a vLAMPS-derived and a PANIS-derived secreted factors were significantly altered under EMS, as compared to NF conditions, mechanistically connecting MASLD and T2DM associated hepatic factors with islet-derived GLP-1 synthesis and regulation. Since vLAMPS-PANIS is compatible with patient–specific iPSCs, this platform represents an important step towards addressing patient heterogeneity, identifying complex disease mechanisms, and advancing precision medicine.

## Introduction

Over 30 million Americans have type 2 diabetes mellitus (T2DM), constituting about 9.4% of the adult population, and about 84 million additional adult Americans have pre-diabetes^1^, amounting to an economic cost of $322 billion annually^2^. Therefore, there is an unmet need for a better mechanistic understanding of T2DM pathophysiology coupled to the development of efficacious and safe treatments to improve patient outcomes. The underlying cause of all forms of diabetes is an inadequate insulin secretion relative to the metabolic needs. The pathogenesis of T2DM is complex and heterogeneous involving multiple tissues with preceding insulin resistance, in the liver, white adipose tissue, skeletal muscle and pancreatic β-cells^3^. Evidence from human and other model organism studies have demonstrated the critical importance of organ crosstalk including the role of myokines^4–9^, adipokines^9–14^, hepatokines^15–19^ and cytokines from immune cells^20,21^ . In parallel with the increase in the incidence of T2DM, there is over a 70% comorbidity between T2DM and metabolic-dysfunction associated steatotic liver disease (MASLD), and it has been determined that patients with MASLD exhibit a higher risk of T2DM than patients without MASLD, suggesting a causal relationship between MASLD and T2DM^22,23^ in some patients. There is a complex relationship between MASLD and T2DM, and evidence demonstrates that MASLD yields an approximate 2-fold increased risk of the development of T2DM independent of obesity and other metabolic risks^24^. While there are several approved and moderately valuable therapies for T2DM, currently the Food and Drug Administration (FDA) has approved only one drug (resmetirom; Rezdiffra^TM^) for the treatment of MASLD^25^. However, resmetirom and many previously failed MASLD drugs have exhibited only up to 25% efficacy in general patient populations^25^. It has been suggested that this is due to the complex pathogenesis of T2DM and MASLD involving factors including genetics, environment, and lifestyle resulting in inter-patient heterogeneity^24,26^. Therefore, a precision medicine approach is required for addressing MASLD-T2DM complexity and heterogeneity.

T2DM is characterized by fasting hyperglycemia and an inability to clear the glucose load that stems from an inadequate insulin release from the pancreatic islet β-cells and inappropriately high glucagon release from the pancreatic islet α-cells. This leads to an increase in gluconeogenesis and *de novo* lipogenesis from the increasingly insulin resistant and steatotic hepatocytes in the liver. The progression of MASLD leads to increased production of several hepatokines, that worsen insulin resistance^27–30^ and, along with the resulting hyperglycemia and increased serum free fatty acids (FFA), cause glucolipotoxicity that worsens β-cell function, in a vicious cycle. Thus, crosstalk between the liver and the islets is critical to the onset and perpetuation of T2DM.

MASLD and T2DM patients exhibit extensive heterogeneity in genotypes and phenotypes that includes the expression of genes that are linked to accelerated disease progression and severity, as well as environment and lifestyle characteristics that also impact disease progression and severity^24^. Current *in vitro* models often do not fully recapitulate critical aspects of complex and heterogenous diseases, and interspecies differences between human and model organism physiology, limit translatability translatability from model organism to humans. To date, many rodent models have been developed and tested but do not reflect the heterogeneity of either MASLD or T2DM, or the combination^31–34^. Simple human cell *in vitro* models do not recapitulate the complex disease due to the absence of important cell-cell and cell-matrix connections^35^. Therefore, it is necessary to implement more complex *in vitro* systems to study human physiology and pathophysiology, as well as disease progression that allow for the study of organ crosstalk and interaction. The development of human liver^36–42^ pancreatic islets^43–47^, and coupled^48–51^ biomimetic microphysiology systems (MPS) that can reflect the genomic, as well as environment and lifestyle heterogeneity offer an important opportunity to recapitulate the heterogeneity of disease.

The use of a combined vascularized liver acinus MPS (vLAMPS)^36,52^ and pancreatic islet MPS (PANIS)^47^ could have an enormous impact on recapitulating many aspects of the liver-pancreatic islet axis. In addition to known insulin resistant liver-derived effectors that can induce β-cell hyperplasia^26,30,53^, we hypothesize that there exists a complementary set of factors derived from the progression to an insulin resistant liver that can potentiate β-cell dysfunction in the context of hyperglycemia and hyperinsulinemia. In this study, we tested our hypothesis by comparing results between stand-alone and physically coupled primary cell-based vLAMPS and PANIS to demonstrate the role of liver-factors influencing pancreatic islets dysfunction. We have harnessed cells from patient-derived cryopreserved cell lots for the liver and cadaver pancreatic islets. We recapitulated the environment and lifestyle phenotypes by progressing the models with media designed to mimic clinical blood chemistries starting with normal fasting (NF) and early metabolic syndrome (EMS) to drive early stage MASLD that can be an optimal starting point for testing therapeutics for early stage. The use of primary cells is important to first establish an adult model system. In parallel, we are developing a patient-derived model based on iPSCs that are differentiated into key liver and pancreatic islets cells that will be used in applications of precision medicine. We further demonstrated the importance of quantifying the reproducibility of the model system so that clinically relevant variations (heterogeneity) can be properly identified, including responses to therapeutics^54–57^.

## Results

To address the hypothesis that the early stage MASLD liver has the potential to directly induce pancreatic islet dysfunction, a modular human structured biomimetic of the liver-pancreas axis was designed and implemented (**Fig. 1**). The overall approach involves a comparative functional characterization of the standalone hepatic (vLAMPS)^36^ and islet (PANIS)^47^ modules in relation to the physically coupled MPS (vLAMPS-PANIS) under both normal fasting (NF) and early metabolic syndrome (EMS) conditions (**Table S1**) using several diverse clinically relevant and reproducible metrics^58^ (**Table 1**).

**Fig. 1.**
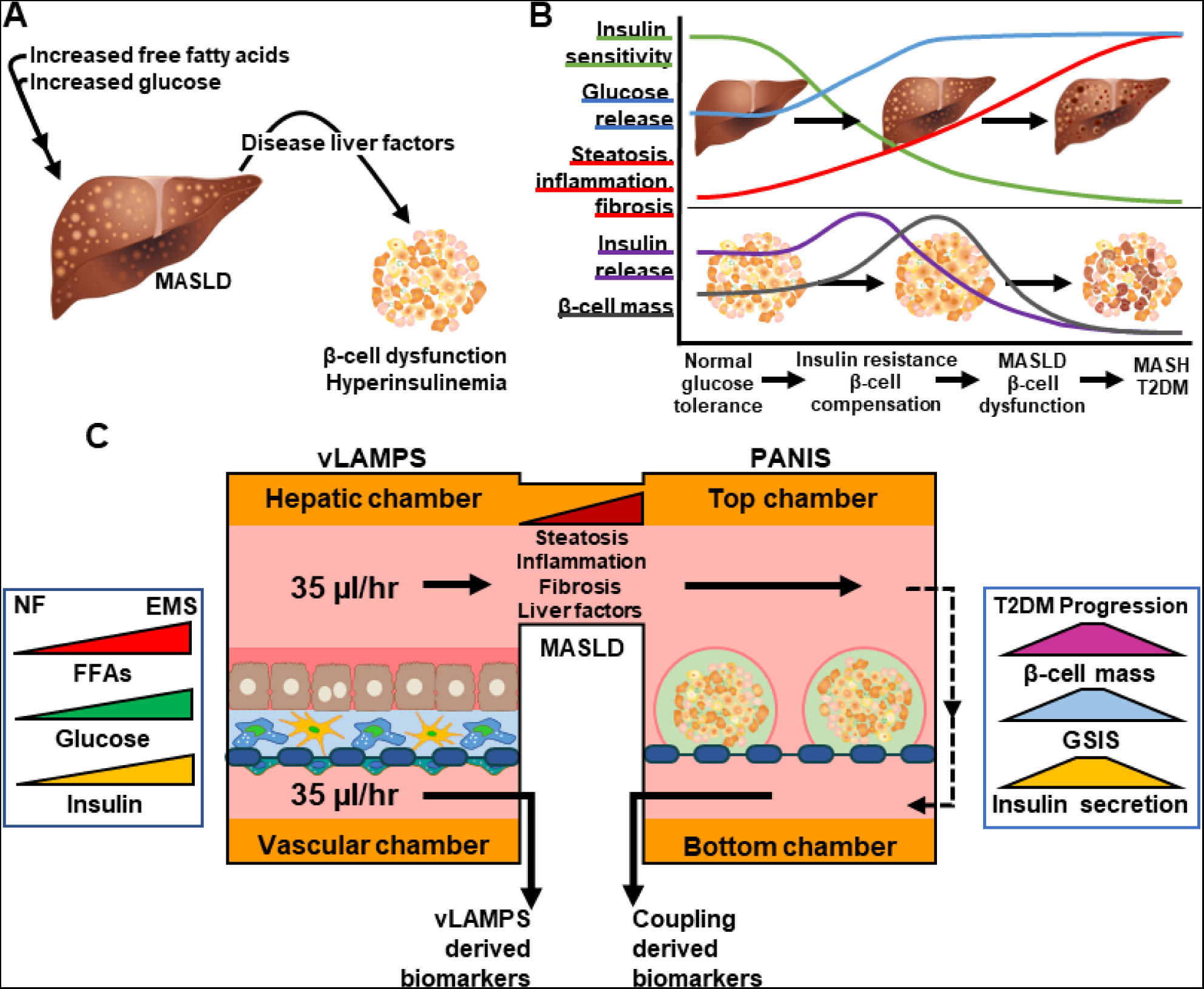
A human liver acinus structured biomimetic MPS coupled to a pancreatic islet MPS recapitulates clinically relevant aspects of the dysregulated liver-pancreas axis associated with the progression of MASLD and T2DM. (**A**) Hypothesis: the early metabolic syndrome (EMS) medium-induced MASLD secretome directly promotes early islet dysfunction. (**B**) Dynamic metabolic dyshomeostasis of the liver and pancreatic islets associated with disease progression. **(C)** Schematic (see Fig. **S1C** for actual photograph) depicting the modular two-organ coupled vLAMPS-PANIS chip. The hepatic and islet modules can also be used in an uncoupled standalone configuration (Fig. **S1A-B**) to enable both intra- and inter-organ functional characterization to be deconvolved under normal (NF) and disease (EMS) conditions. The hepatic influx medium composition determines a normal fasting (NF) or EMS state and its continuous flow among the self-assembled multiple cell types (see **Methods and Results**) can be monitored for biomarkers that complement in situ measurements. Statistical comparisons

**Table 1.**
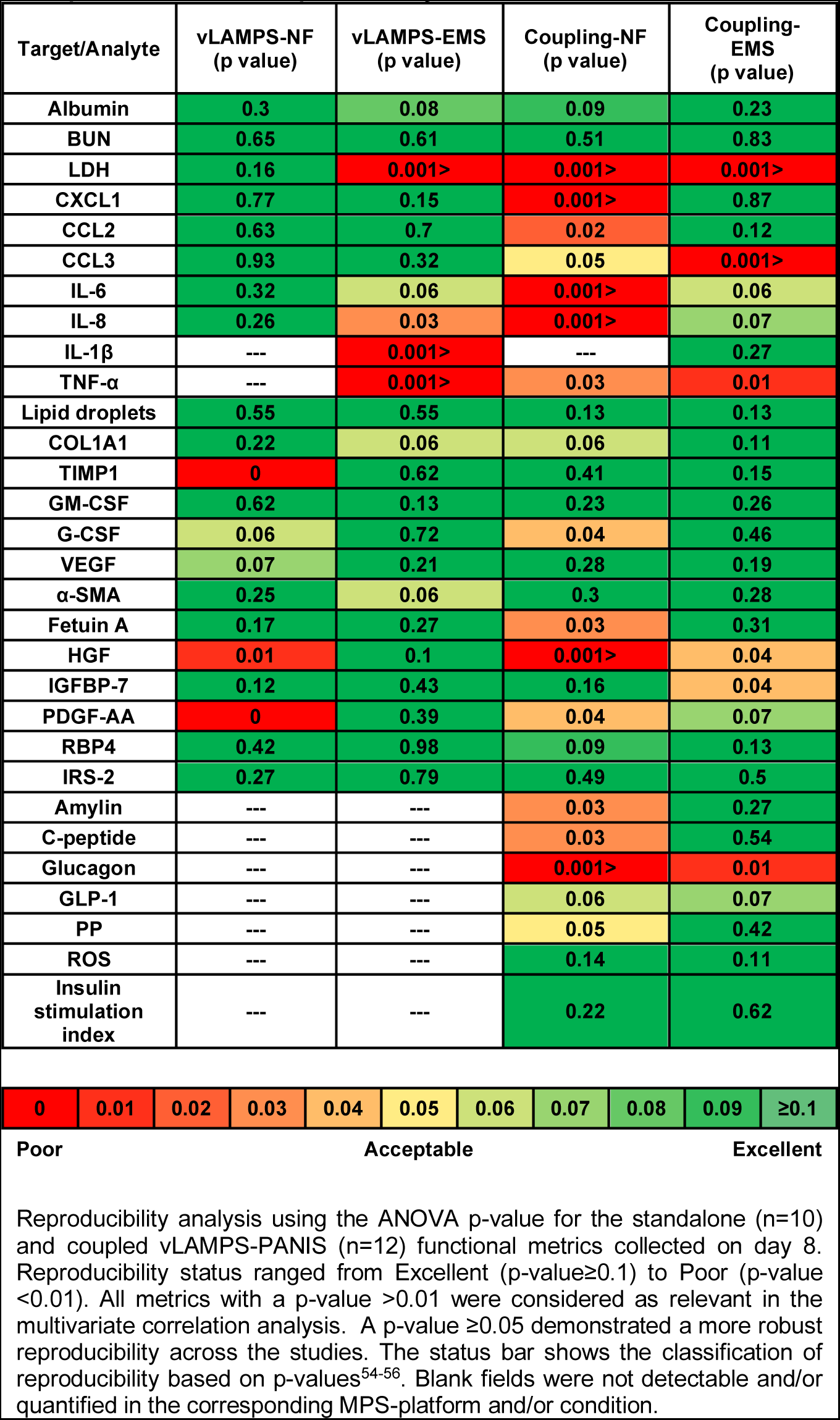
Several clinically relevant metrics from standalone vLAMPS and coupled vLAMPS-PANIS, critical for the interpretation of the study results, exhibit acceptable to excellent reproducibility.

### Characterization of standalone vLAMPS and PANIS modules

The assembly of the vLAMPS with primary human hepatocytes and liver sinusoidal endothelial cells (LSECs), and human cell lines for stellate and Kupffer-like cells, which is composed of a vascular channel separated from a hepatic channel (chamber) by a porous membrane that allows communication between these channels, recapitulates the 3D structure of the liver acinus^36^ **(Fig. S1A)**. Structural features within the vLAMPS that could impact the composition of the hepatic secretome include the space of Disse that mediates non autonomous cell signaling involving in part the LECM^36,59^ (**Fig. S1A**) and the continuous oxygen zonation ranging from 16%-5% over three zones (**Fig. S2A-E**) that modulates oxidative phosphorylation, metabolism, and gene expression^60^. The oxygen zonation of the vLAMPS was first modeled (**Fig. S2A**) and then experimentally validated with oxygen sensitive beads^36^ (**Fig. S2B-E**) in the context of a uniform distribution of hepatocytes and LSECs (**Fig. S2E** and **S2F**) whose oxygen consumption creates the gradient. In addition, we show that steatosis increases across the zones, consistent with the physiologically elevated *de novo* lipogenesis in zone 3^58,61^ (**Fig. S2H and S2I**). Basic hepatocyte function assessed by albumin and urea synthesis and secretion is evident under both NF and EMS conditions (**Fig. S3A and B**) over the 8-day time course, and no appreciable progressive plasma membrane damage is observed as determined by a LDH release assay (**Fig. S3C**). The NF and EMS media conditions were previously shown to recapitulate normal fasting and early stage MASLD in LAMPS (liver acinus microphysiology system) using a panel of clinically relevant MASLD progression metrics that includes hepatic steatosis, immune activation, stellate cell activation, and the production of pro-fibrotic markers^58^. We now show in vLAMPS that EMS induces early stage MASLD using a more comprehensive panel of metrics. Relative to perfusion under NF conditions, perfusion with EMS in the standalone vLAMPS showed an increase in steatosis (**Fig. 2A and 2B**) and a decrease in IRS-2 expression (**Fig. 2C and 2D)** on day 8, indicative of impaired insulin signaling^62^. However, no significant difference in α-SMA expression (**Fig. 2E and 2F**) by immunofluorescence image analysis was observed suggesting that activation of quiescent stellate cells to proliferative fibrogenic myofibroblasts was not fully induced by EMS perfusion. This is consistent with no significant increase in interstitial collagen (COL1A1) in the secretome (**Fig. 2F**), although the overall trend in COL1A1 secretion shows an increasing trend in EMS medium (**Fig. S4**). IL-6 (**Fig. 2B**) and GM-CSF (**Fig. 2F**) showed significant increases under EMS perfusion compared to NF perfusion, indicating progression towards the pro-inflammatory state. As expected, the islet-specific secreted factors, amylin, C-peptide, glucagon, GLP-1, and pancreatic polypeptide, were not detected in the standalone vLAMPS secretome (**Fig. S4**). Together these results suggest that an eight-day EMS perfusion in the standalone vLAMPS results in steatosis, impaired insulin signaling, and an altered proinflammatory secretome with no appreciable fibrogenesis, consistent with the early stages of MASLD progression. These metrics were reproducible across independent studies (**Table 1**) enabling biological differences at different times and/or under different perfusion conditions to be discerned.

**Fig. 2.**
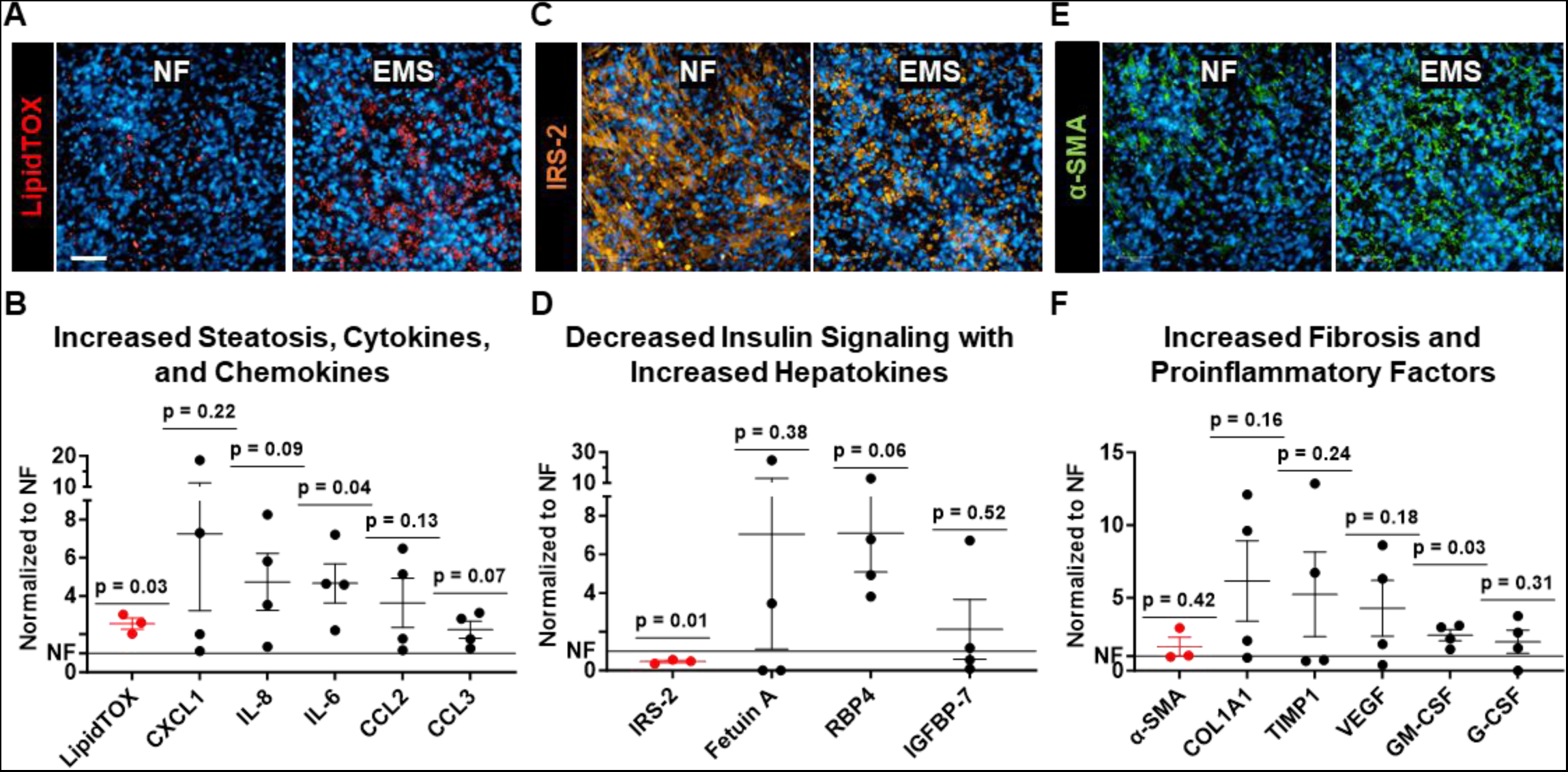
EMS medium promotes an early MASLD phenotype in the standalone vLAMPS maintained for 8 days. (**A**) Representative immunofluorescence images of lipid accumulation (LipidTOX) in hepatocytes under NF and EMS conditions and (**B**) LipidTOX quantification in conjunction with secreted proinflammatory cytokines and chemokines. (**C**) Representative immunofluorescence images of Insulin receptor susbtrate-2 (IRS-2) in hepatocytes under NF and EMS conditions and **(D)** IRS-2 quantification in conjunction with an increase in hepatokine secretion. (**E**) Representative immunofluorescence images of stellate cell activation (α-SMA) under NF and EMS conditions and (**F**) α-SMA quantification in conjunction with secreted fibrotic and proinflammatory factors. Scale=100µm. Scatter plot graphs display the mean and standard error (SEM) of the EMS values normalized to NF for each of the metrics. (see Fig. **S4**). Points on the graph represent individual chips. Statistical analysis by Wilcoxon signed-rank test of normalized EMS values to their corresponding NF study.

The seeding of cadaver-derived islets onto the porous PET membrane (**Fig. 1C and S1B**) and encapsulation within polymerized alginate hydrogel to maintain their 3-dimensional structure and function over a 2-week period in the assembled PANIS have been described^47^ and detailed in the **Methods**. We compared the ability of the chemically defined NF in PANIS to maintain islet phenotype and function with the proprietary Prodo Labs medium PIM(R) (**Fig. S5**). No significant difference between viability of C-peptide or glucagon expressing cells was observed (**Fig. S5D**). The ratio of insulin secreted per islet at 16mM to 3mM glucose is the insulin stimulation index (ISI), which is a critical test of islet function. The amount of insulin secreted per islet was higher in NF versus PIM(R) under both low and high glucose conditions (**Fig. S5E**), however no significant difference in the insulin stimulation index between the two media (**Fig. S5F**) was apparent. We then compared the structure and function of the islets in the standalone PANIS under NF and EMS conditions (**Fig. 3**). There was no significant difference in cell viability between NF and EMS media (**Fig. 3A** and **3B**). Consistent with this result, immunofluorescence imaging showed no difference between the two conditions in the relative fraction of C-peptide expressing β-cells to glucagon-expressing α-cells (**Fig. 3C** and **3D**). Clinically, ROS is associated pathophysiologically with oxidative stress and physiologically with homeostatic regulation of metabolic processes that include insulin secretion in islets^63,64^. DCFDA-based fluorescence staining for ROS showed no significant difference between the two perfusion conditions (**Fig. 3E and 3F**). Furthermore, no significant differences in the secretion of islet-specific factors were observed between the two conditions (**Fig. 3G and S6).** To evaluate islet insulin response, islets were retrieved on day 8 and analyzed for glucose stimulated insulin secretion (GSIS) under 3mM and 16mM glucose. Under both NF and EMS perfusion conditions, there was a healthy GSIS response (**Fig. 3H**) with only a trend towards a lower insulin stimulation index under EMS versus NF conditions (**Fig. 3I)**. Thus, in contrast to changes induced by EMS compared to NF in the standalone vLAMPS, significant media induced changes in the standalone PANIS were not evident in EMS indicating that perfusion with this medium alone for 8 days was not sufficient to drive islet dysfunction. We next addressed whether the EMS-induced changes in vLAMPS steatosis, insulin signaling and its secretome could promote islet dysfunction in the physically coupled vLAMPS-PANIS.

**Fig. 3.**
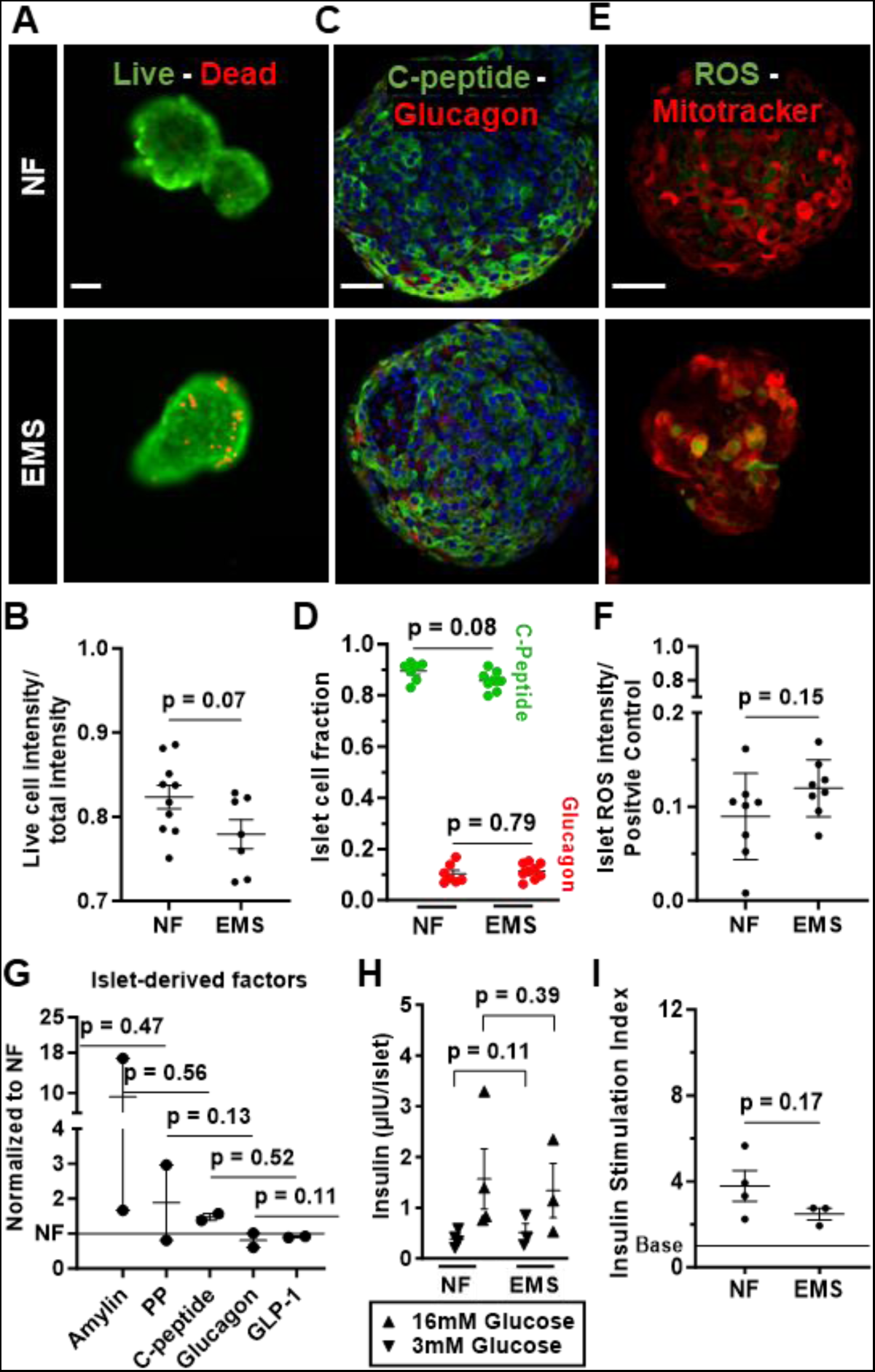
EMS medium alone is not sufficient to induce islet dysfunction. NF and EMS media maintain islet viability and functionality including glucose stimulated insulin response for 8 days in the standalone PANIS. (**A**) Representative viability (live/dead) image for islets cultured in NF and EMS media. (**B**) Viability quantification shows no significant difference between NF and EMS. (**C**) Immunofluorescence labelling for C-peptide expressing β-cells and glucagon expressing α-cells for islets cultured in NF and EMS. (**D**) The fraction of quantified C-peptide and glucagon positive cells shows no significant difference. (**E**) Representative images showing mitochondrial associated ROS immunofluorescence labelling of islets. (**F**) Percentage of ROS intensity quantification with respect to a positive control (50 µM TBHP) for islets cultured in NF and EMS media (see **Methods**) demonstrates no significant difference between NF and EMS. (**G**) Standalone PANIS efflux analysis showed the change in islet-derived factors in EMS normalized to NF (see **Fig. S6**), with no significant difference between them. (**H**) Glucose stimulated insulin secretion (GSIS) assay, where islet samples were subjected to low (3mM), and subsequent high (16mM) glucose concentration and the resulting secreted insulin was measured and showed no significant difference at either glucose concentration in islets that were maintained in either NF or EMS media. (**I**) Comparison of insulin stimulation index secretion for islets cultured in NF and EMS showing no statistically significant difference. Scatter plot graphs display the mean and standard error (SEM) with p-values shown. For graphs -*B, D, F, H*, and *I*-statistical analysis was done by two-tail t test with Welch’s correction. For graph -*G*-Statistical analysis was done by Wilcoxon signed-rank test of normalized EMS values to their corresponding NF study. All scale=50µm.

### Characterization of the coupled vLAMPS-PANIS

The physical coupling of vLAMPS and PANIS is described in the **Methods** and configured with the hepatic chamber efflux of the vLAMPS serving as the influx for the PANIS (**Fig. 1C, S1C**). Similar to the standalone vLAMPS, the basic hepatocyte function of albumin and urea synthesis and secretion is evident under both NF and EMS conditions (**Fig. S7A** and **S7B**). Over the 8-day time course, no appreciable progressive plasma membrane damage is observed as determined by an LDH release assay (**Fig. S7C**). Analysis of diverse secretome measurements across independent studies for both NF and EMS conditions showed overall acceptable to excellent reproducibility of the metrics (**Table 1**) enabling biological differences at different times and/or under different perfusion conditions to be discerned. Consistent with the results from the standalone vLAMPS, an increase in steatosis (LipidTOX) and a decrease in IRS-2 expression by EMS relative to perfusion with NF are seen (**Fig. 4A** and **4B**), with the latter indicative of impaired insulin signaling being induced in the vLAMPS module. Consistent with **Figure 2**, no significant increase in α-SMA was evident; however, an EMS induced increase in secreted COL1A1 in the coupled efflux suggested a trend towards systemic fibrogenesis, consistent with the early stages of MASLD progression (**Fig. 4A** and **4B**). Analogous to the standalone PANIS, the viability of the islet cells remained constant between NF and EMS and no significant change in the relative distribution of β- to α-cells suggesting that in the coupled vLAMPS-PANIS the islets were structurally intact with viable cells under both perfusion conditions (**Fig. 4C-E**). Although there was no significant difference in ROS levels between NF and EMS conditions (**Fig. 4F**), as observed in the standalone PANIS (**Fig. 3F**), the levels of ROS were 2-fold higher in the coupled system suggesting that hepatic factors from the vLAMPS module increased ROS levels in the PANIS module (**Fig. S8**). Coupling potentiated an EMS-dependent significant increase in the levels of the islet-specific secreted factors, amylin and pancreatic polypeptide (**Fig. 4G**). Under NF conditions, a relatively lower secretion of insulin per islet was observed at 3 mM glucose compared to EMS, and under EMS conditions a trend toward lower insulin was observed at 16 mM glucose compared to NF (**Fig. 4H**). In contrast to the standalone PANIS (**Fig. 3H** and **3I**), the coupled system exhibited a highly significant reduction in glucose stimulated insulin secretion index under EMS versus NF perfusion conditions (**Fig. 4I).** These results are consistent with the hypothesis that a healthy liver under NF conditions may impart homeostatic islet regulation to prevent hypoglycemia whereas a steatotic MASLD liver with impaired insulin signaling may predispose the islets towards a diminished capacity to secrete insulin at higher glucose. Corroborating these observations under NF conditions and consistent with this hypothesis, C-peptide secretion in the coupled vLAMPS-PANIS under NF conditions was two-fold lower than its secretion in the standalone PANIS while no difference due to coupling under EMS conditions was observed (compare **Fig. S6 and S9**).

**Fig. 4.**
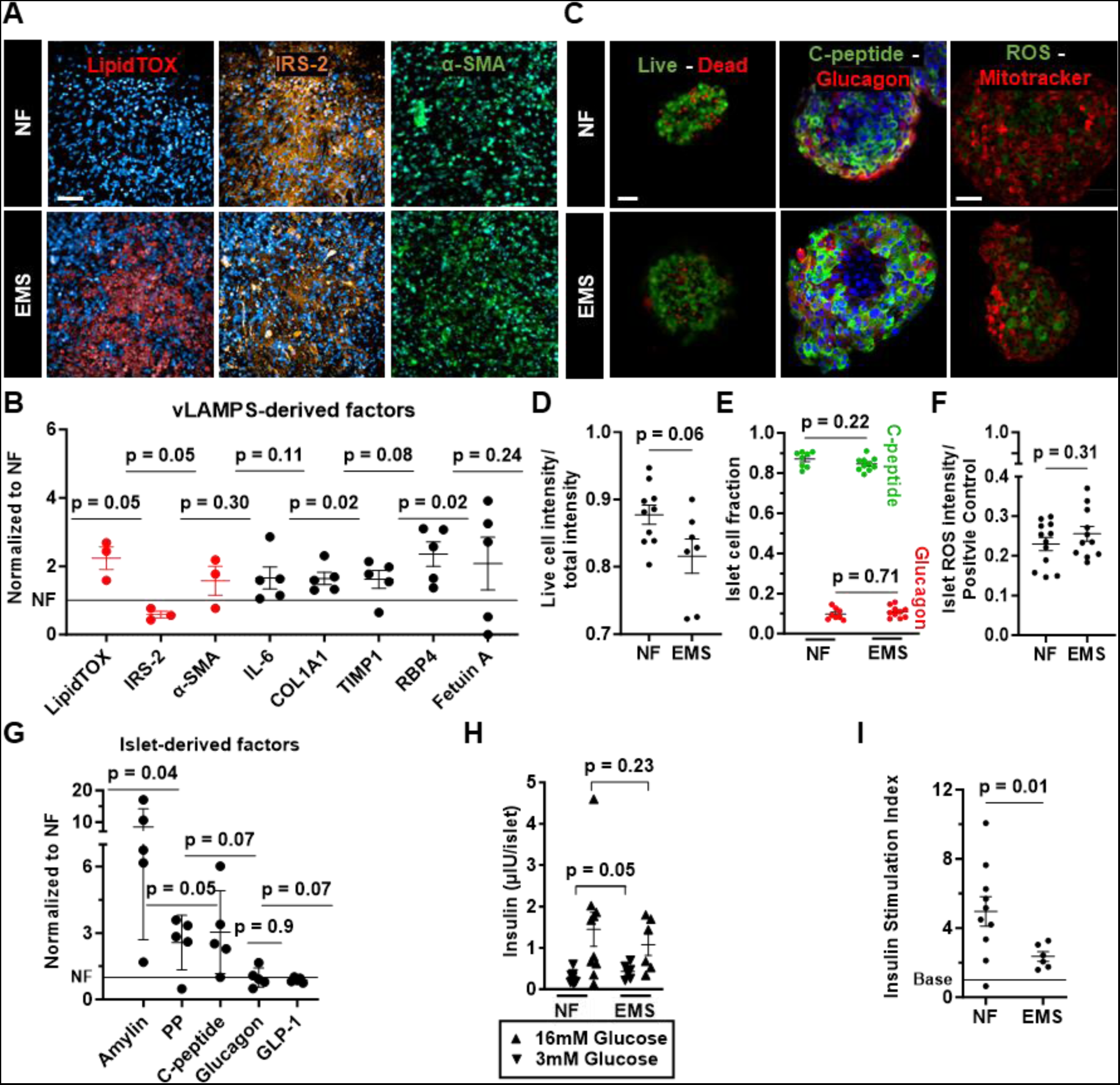
Coupling vLAMPS to PANIS maintains islet viability under EMS conditions for at least 8 days but induces significant perturbations in the islet secretome and function. (**A**) Representative images demonstrating the effect of EMS medium on vLAMPS indicated by markers of steatosis (LipidTOX), insulin signaling (IRS-2), and fibrosis (α-SMA) in hepatocytes. Scale=100µm. (**B**) Corresponding quantitation of hepatic EMS induced steatosis, decreased insulin signaling, and increasing fibrosis (red). Along with vLAMPS-derived EMS secreted factors normalized to complementary NF chips (reproducible, **Table 1**). (**C**) Representative images of viability (live/dead), immunofluorescence labeling for C-peptide (green) expressing β-cells and glucagon (red) expressing α-cells, and ROS oxidative stress for coupled islets cultured in NF and EMS. All scale=50µm. (**D**) Viability quantification shows no statical difference between conditions. (**E**) No statistically significant difference between the fraction of quantified C-peptide and glucagon positive cells. (**F**) Percentage of ROS intensity quantification with respect to a positive control for coupled islets cultured in NF and EMS (see **Methods**) demonstrated no significant differences between islets cultured in either NF or EMS media. (**G**) islet-derived factors in EMS normalized to NF (reproducible, **Table 1**) demonstrated significant increases in the secretion of both amylin and pancreatic polypeptide in EMS medium. (**H**) The islets from the coupled vLAMPS-PANIS in EMS exhibit significant abnormal insulin response to low glucose in islets that were maintained in either NF or EMS media, which was not observed in the standalone PANIS islets (see **Fig. 3H**). (**I**) The same significance is observed in the insulin stimulation index response of islets that were coupled with NF medium compared to EMS medium. (see Methods). Scatter plot graphs display the mean and standard error (SEM) with p-values shown. For graphs -*D, E, F, H,* and *I*-statistical analysis was done by two-tail t test with Welch’s correction. For graph -*B* and *G*-statistical analysis was done by Wilcoxon signed-rank test of normalized EMS values to their corresponding NF study.

### Correlation analysis supports functional response in coupled vLAMPS-PANIS

To gain further evidence and insights for communication between the two modules in the coupled vLAMPS-PANIS and test the hypothesis that liver derived factors can modulate islet derived factors important for T2DM progression, we assessed pairwise correlations between secreted factors selectively derived from the vLAMPS module and factors selectively derived from the PANIS module under NF and EMS conditions (defined as a pair). The reproducibility for the measurement of each secreted factor is presented in **Table 1**^56^. Correlation analysis was restricted to hepatic and islet factors with reproducibility ANOVA p-values ≥ 0.01 (**see Table 1**). Statistically significant correlation coefficients were observed in at least one medium condition for 14 of the 23 reproducible pairs (correlation coefficient p-value ≤ 0.05, **Table 2**). Six of these 14 pairs (marked with an asterisk in **Table 2**) were between hepatic and islet factors with a more stringent reproducibility ANOVA p-value ≥ 0.05 in both NF and EMS conditions. As expected, the 3 statistically significant correlation coefficients derived from these six pairs (Table 2) were also identified in the less stringent analysis along with 13 other statistically significant correlation coefficients derived from the total of 14 pairs (as discussed below). This result indicated that amid potentially lower reproducibility of single measurements using the less stringent reproducibility criterion, statistically significant covariances could still be identified.

**Table 2.** Pairwise correlations involving a vLAMPS derived factor and an islet derived factor in the coupled vLAMPS-PANIS efflux are altered under NF versus EMS conditions consistent with a role of hepatic factors in the dysregulation of islet function.

| vLAMPS-derived | Islet-derived | NF Correlation coefficient (p-value) | EMS Correlation coefficient (p-value) |
| --- | --- | --- | --- |
| TNF-α | Amylin | 0.78 (0.04) | 0.82 (0.09) |
|  | C-peptide | 0.70 (0.08) | 0.59 (0.29) |
|  | GLP-1 | -0.80 (0.03) | -0.78 (0.12) |
|  | PP | 0.63 (0.13) | 0.73 (0.16) |
| COL1A1 | Amylin | 0.81 (0.03) | 0.80 (0.10) |
|  | C-peptide | 0.99 (<0.001) | 0.58 (0.30) |
|  | GLP-1* | -0.24 (0.61) | -0.62 (0.26) |
|  | PP* | 0.83 (0.02) | 0.89 (0.04) |
| TIMP1 | GLP-1* | 0.71 (0.07) | 0.29 (0.63) |
| GM-CSF | GLP-1* | 0.77 (0.04) | 0.62 (0.27) |
| G-CSF | Amylin | 0.79 (0.03) | 0.71 (0.18) |
|  | C-peptide | 0.98 (<0.001) | 0.49 (0.40) |
|  | GLP-1 | -0.20 (0.67) | -0.68 (0.21) |
|  | PP | 0.84 (0.02) | 0.68 (0.21) |
| Fetuin A | Amylin | 0.81 (0.03) | 0.17 (0.78) |
|  | C-peptide | 0.92 (<0.001) | -0.06 (0.92) |
|  | GLP-1 | -0.13 (0.78) | -0.80 (0.10) |
|  | PP | 0.69 (0.09) | -0.10 (0.87) |
| IGFBP-7 | GLP-1 | 0.20 (0.66) | 0.65 (0.24) |
| RBP4 | Amylin | 0.76 (0.05) | 0.89 (0.04) |
|  | C-peptide | 0.81 (0.03) | 0.65 (0.24) |
|  | GLP-1* | -0.03 (0.95) | -0.76 (0.14) |
|  | PP* | 0.59 (0.16) | 0.87 (0.05) |

Using the p>0.01 reproducibility threshold, 23 secreted factors pairs (one derived primarily from the vLAMPS and the other from PANIS) were identified and their correlation coefficients determined under both NF and EMS conditions (**Table 2**). Focusing exclusively on the statistically significant correlations (p≤0.05) in at least one medium condition there were 8 pairs that had at least a 20% change between two conditions (highlighted in blue), and 6 that did not change (shown in gray) (**Table 2**). Seven of these 8 pairs showed a significant correlation only under NF conditions. Conversely one correlation involving RBP4 and pancreatic polypepetide was only significant under EMS conditions. Particularly striking was the involvement of C-peptide (a direct marker of newly processed and secreted insulin) in 4 of these 7 correlations and 1 more involving the vLAMPS-derived cytokine, GM-CSF, and islet-derived GLP-1, that promotes insulin secretion. The loss of these 5 positive correlations under EMS conditions is consistent with the EMS-induced dysregulation of GSIS (**Fig. 4I**). In addition to their positive correlations with C-peptide under NF conditions, Fetuin A and RBP4 showed a strong trend toward a negative correlation with GLP-1 under EMS conditions (**Table 2**). This observation is complementary to the aforementioned positive correlations and further supports a sensitive condition-dependent liver-pancreas signaling network. These statistical based correlations can guide us toward a mechanistic model supported by the literature and are discussed below.

## Discussion

The results presented in this study show that vLAMPS-PANIS can serve as a human biomimetic recapitulating critical aspects of the early stages of MASLD development, including the development of insulin resistance, leading to pancreatic β-cell dysfunction. Its modular design can be used to distinguish inter-organ from intra-organ signaling and help to deconvolve the pleiotropic effects of secreted molecules. Its implementation here has provided evidence supporting the hypothesis that an early stage MASLD liver can induce β-cell dysfunction through a hepatic-pancreatic axis that at least in part underlies the high comorbidity among MASLD and T2DM patients^23^. The diverse panel of metrics that is reproducible across independent experiments has 1) enabled pairwise correlations between a vLAMPS and a PANIS secreted factor to be identified and 2) their differential dependence between normal fasting and early metabolic syndrome conditions to be determined, lending further support for this working hypothesis.

These correlations and their dependencies on media-induced conditions provide not only statistical-based support for the liver-pancreas axis but in conjunction with clinical, *in vitro*, and animal model data give rise to mechanistic models and causal hypotheses. Fetuin A, a hepatokine associated with MASLD and T2DM^65–67^, is exclusively secreted from the vLAMPS module of vLAMPS-PANIS and under NF conditions is positively correlated with C-peptide, the direct and specific marker of newly processed and secreted insulin. A similar relationship between Fetuin A and insulin has been observed in animal models involving Fetuin A knockout studies and its exogenous administration^68,69^ and in patients^70^, and results from Fetuin A binding to hepatic insulin receptor, inhibiting its receptor tyrosine kinase (RTK) activity to induce a homeostatically regulated rise in insulin levels (i.e., C-peptide)^68,71^. Under EMS conditions a positive correlation between Fetuin A and C-peptide is no longer observed in the vLAMPS-PANIS. Mechanistically, circulating Fetuin A in the context of free fatty acids can bind directly to β-cells through the TLR-4, promoting a NF-kB dependent inflammatory response that disrupts glucose stimulated insulin secretion (GSIS)^28,72^. Consistent with this mechanism, a negative correlation between Fetuin A and GLP-1, the hormone that promotes GSIS, was observed only under EMS conditions that create the MASLD disease state in the liver. In studies involving mouse islets, Fetuin A in the context of free fatty acids and working through the TLR-4-NF-kB pathway was shown to enhance the expression of DPP-IV, the dipeptidyl peptidase responsible for degrading GLP-1^73^. A complementary mechanism for the observed negative correlation between Fetuin A and GLP-1 likely involves the fetuin-dependent diminished capacity of inflamed β-cells to conduct the bidirectional paracrine signaling with α-cells that is required for GLP-1 processing and secretion^74,75^. By analogy, the retinol binding protein RBP4 that is associated with MASLD and T2DM^76^ and is secreted predominantly from vLAMPS under EMS conditions, has a negative correlation with GLP-1 under these conditions. As a complex with retinol, it has been shown in mouse systems to induce direct β-cell dysfunction through the RBP4/STRA6/JAK2/STAT1/insulin gene enhancer protein 1 to lower GSIS and insulin synthesis^77,78^. Together, this mechanistic analysis of the statistical correlations derived from vLAMPS-PANIS under EMS and NF conditions supports a model in which clinically relevant factors secreted from early stage MASLD liver can directly interact with islet β-cells to perturb the finely tuned paracrine signaling critical for GLP-1 regulation of GSIS^74^ (Supplementary **Fig. S11**). These liver secreted factors, as exemplified by Fetuin A, are multifaceted. Under NF conditions they can act in an autocrine or paracrine manner as normal homeostatic components of glucose regulation by modulating hepatic insulin signaling. Under EMS conditions these factors either become overexpressed or in the presence of FFAs assume a pathophysiological endocrine role to dysregulate intra-islet signaling. The functional, statistical and mechanistic analysis of this vLAMPS-PANIS study support the commonality of this mechanism to engender MASLD-T2DM comorbidity.

This current two-organ modular version of vLAMPS-PANIS does not include intestine-derived GLP-1^79^ indicating the potential importance of pancreatic islet-derived GLP-1 in normal homeostasis and its dysregulation in early metabolic syndrome disease. Although this study focused on early-stage metabolic syndrome disease, positive correlations between the later stage liver stellate cell-derived fibrogenesis marker, COL1A1, and islet derived amylin and pancreatic polypeptide were observed under EMS conditions. This may represent the beginning of centrally mediated compensatory mechanisms involving amylin and pancreatic polypeptide that regulate satiety and gastric emptying when GSIS is compromised^80^. The correlation between stellate cell derived COL1A1 and islet F-cell derived pancreatic polypepetide^81^ indicates the capability of vLAMPS-PANIS to detect indirect but potentially important signaling interactions between relatively rare cell populations. Furthermore, in addition to the correlations made for NF and EMS media, we have also previously demonstrated the use of late metabolic syndrome (LMS) medium in liver MPS^58^. This medium contains pro-fibrotic molecular drivers including TGF-β and lipopolysaccharide (LPS), which recapitulate features in late stage MASLD^58^. Although not performed in the current study, correlations between hepatic and pancreatic factors using this medium will provide additional mechanistic insights in the future. However, here we focused on the earlier stage in the MASLD-T2D progression as a reasonable starting point to explore mechanisms and therapeutic strategies.

To determine the clinical relevance of vLAMPS-PANIS, well established markers implicated in MASLD and T2DM were employed. Although this supervised approach enabled this objective and provided evidence supporting the underlying role of the liver-pancreas axis in MASLD and T2DM comorbidity, it nevertheless limited the full potential of vLAMPS-PANIS to provide a holistic understanding of physiological and pathophysiological mechanisms. We anticipate that future studies employing vLAMPS-PANIS will take a more comprehensive unbiased approach that utilize larger panels of metrics. These panels will include multiple ‘omics’ analyzed at different time points complemented by real-time imaging under media conditions that mimic both early and late-stage metabolic syndrome disease. Since MASLD and T2DM involve multiple organs in addition to the liver and the pancreas, we expect that future studies will exploit the modular design of vLAMPS-PANIS to include the coupling of human microphysiology systems comprising adipose tissue, the intestine, and/or gut microbiome. As demonstrated with the coupled vLAMPS-PANIS in this study, the capability of systematically constructing and deconstructing complex disease biomimetic models provides a powerful approach for deciphering pathophysiology and identifying targetable mechanisms.

The use of primary cells was important to establish an MPS disease model that reflected fully matured cells. This will be critical in interpreting the use of patient-based, induced pluripotent stem cell (iPSC)-derived cells for the liver and pancreatic islets. However, the use of primary cells has the limitations of supply of the same cells for multiple studies over time and the ability to generate patient-derived or patient-specific MPS to define heterogeneity. The use of optimally matured, patient-derived iPSCs and synthetic biology in the disease MPS addresses the limitations of primary cells and allows the application of precision medicine to understand inter-patient heterogeneity that will inform optimal therapeutic strategies for individual patients or patient cohorts with the most similar characteristics.

## Methods

### Human hepatocytes and non-parenchymal cells (NPCs)

Cryopreserved human primary hepatocytes (post-thaw plateability with >90% viability) were purchased from Thermo Fisher (Hu1960). Hepatocyte donor information is available in supplementary **Table S2**. Liver sinusoidal endothelial cells (LSECs) were purchased from Lifenet Health (cat. no. NPC-AD-LEC-P1). To preserve a consistent LSECs stock, cells were thawed and expanded in EGM-2 endothelial cell growth medium (Lonza, cat. no. CC-3124) for two passages (P-2) and re-frozen at 1×10^6^ cells/ml. Human hepatic stellate cell line LX-2 were acquired from Sigma (cat. no. SCC064), and were cultured in DMEM-2 (Thermo Fisher, cat. no. 10564011) supplemented with 2% FBS (Fisher Scientific, NC9525043) and 10mg/ml Pen-Strep (Hyclone, cat. no. SV30010). Human monoblast cell line THP-1 were purchased from ATCC (cat. no. TIB-202). THP-1 cells were used as Kupffer-like cells, as they can be differentiated into activated macrophages^82^. The cells were expanded in RPMI 1640 (Thermo Fisher, cat. no. 11875093) supplemented with 10% FBS, 10mg/ml Pen-Strep, and 2 mM L-Glutamine (Hyclone, cat. no. SH30034.01). Forty-eight hours prior to harvesting for seeding into the vLAMPS, cells were treated with 200 ng/mL phorbol myristate acetate (PMA) (Calbio, cat. no. 524400) in RPMI^83^. Prior to MPS assembly, all cultured cells were maintained at maximum of 80% confluency with their respective cell culture media and incubated at 37 °C in 5% CO2. LX2 and THP-1 cells were kept under 20 passages. Hepatocytes were maintained in liquid nitrogen until thawed to be seeded in the vLAMPS.

### Pancreatic islets

Human cadaveric pancreatic islets were procured from Prodo Labs (San Francisco, CA) and maintained in suspension for four days using the proprietary Prodo islet media for recovery (PIM(R)) (Prodo Labs, cat. no. PIMCR001GMP). Donor details can be found in supplementary **Table S3**. The islets were maintained in a petri dish (Fisher Scientific, cat. no. FB0875711) suspended in 10mL PIM(R), for a minimum of 4 days. Fifty percent medium changes were carried out every other day under static suspension and incubated at 37 °C in 5% CO2.

### Normal fasting and early metabolic syndrome media

The development of normal fasting (NF) and early metabolic syndrome (EMS) media have been previously described^58^ (Supplementary **Table S1**). EMS medium consisted of NF medium with the modification of 20mM glucose (Sigma, cat. no. G8644ML), 30pM glucagon (Sigma, cat. no. G2044), 10nM insulin (Gibco, cat. no. 12585-014), and addition of 200 µM oleic acid-BSA conjugated (Cayman, cat. no. 29557) and 100 µM sodium palmitate-BSA conjugated (Cayman, cat. no. 29558) (Supplementary **Table S1**).

### Microfluidic glass device design

The vLAMPS and PANIS were constructed using the Micronit Organ-on-Chip glass microfluidic platform^36^. The 2-chamber flow device consists of three borosilicate glass layers (Micronit, cat. no. OOC 00739). The middle layer contains an oval orifice the bottom of which is covered by porous PET membrane, creating an oval-well on the top, partitioning the top and the bottom chambers. The vLAMPS has an oval area of 77mm^2^ with 0.4µm pores (Micronit, cat. no. 03237), while the PANIS uses the 100mm^2^ oval with 3µm pores (Micronit, cat. no. 02861). The layers are held together within a 4-device aluminum clamp (Micronit, cat. no. 4515) which has the openings for the corresponding inlets and outlets.

The vLAMPS fluid flow configuration was achieved by parallel perfusion of the 2 chambers denoted hepatic and vascular chambers (Supplementary Fig. **S1A**), with two inlets and two outlets. The PANIS fluid flow configuration consisted of a single inlet/outlet achieved by a looped path through the chip by interconnecting the two chambers (top and bottom chambers) (Supplementary Fig. **S1B**).

### Computational fluid dynamics (CFD) and vLAMPS oxygen tension

A perfusion flow rate was estimated based on computational fluidic dynamics (CFD) to establish a physiological oxygen zonation gradient in vLAMPS. The CFD modeling of the oxygen zonation using a transport of diluted species model and a flow model has been previously described^36^. In the geometry of the current middle layer (oval area and PET membrane) the number of hepatocytes was decreased from published study^36^, but the cell’s oxygen consumption rate was maintained at 0.45 nmol/s/10^6^cells^84,85^. The PET membrane’s porosity, thickness, and oxygen diffusion coefficient were updated accordingly^86^ in the CFD model (Supplementary **Table S4**). The estimated flow rate of 35µl/hr was determined to generate a gradual oxygen tension ranging from 1.8×10^-1^ to 4×10^-1^ mol/m^3^ with ∼85,000 hepatocytes on a parallel perfusion in a glass device.

The flow rate and oxygen tension were empirically validated with the previously described ratiometric imaging of oxygen sensitive beads to reference beads^36^. In the vLAMPS oval area, the regions of interest (ROIs) were arrangements of 12 columns equidistantly distributed across the long axis of the oval. Each column is made up of 4 fields (20x magnification) evenly distributed perpendicular to each side of the center of the oval’s long axis, a total of 48 fields across 12 ROIs.

### vLAMPS fabrication

The fabrication of the vLAMPS was as previously described^36^ with the following modifications. The vLAMPS middle layer was treated with a solution of 100 μg/mL fibronectin (Sigma, cat. no. F1141) and 150 μg/mL collagen (Corning, cat. no. 354249) in 1x phosphate buffered saline (PBS) (Gibco, cat. no. 21600-069) overnight in a 4°C cold room. Based on the proportions of liver cell types *in vivo* seeding numbers were determined by allometric scaling to be 40: 14: 6, Hepatocytes: Kupffer: Stellate^37,87^. LSEC seeding ratio is 1:1 to hepatocytes to cover the extended surface area of the PET membrane as previously described^36^. The cell densities used here were: P-2 LSECs 8.5×10^5^ cells/ml, LX2 at 1.3×10^4^ cells/ml, PMA-treated THP-1 at 2.4×10^4^ cells/ml, and hepatocytes at 2.12×10^6^ cells/ml. The decellularized porcine liver extracellular matrix (LECM) was provided by Dr. Badylak^88^. To promote hepatocyte polarization, an overlay of rat tail collagen type 1 solution (2.5 mg/mL) in NF medium (or EMS for disease conditions) was deposited on top of the hepatocytes inside the borders of the oval-well. Lastly, the middle layer was fully submerged in NF or EMS medium and kept at 37°C and 5% CO_2_ until assembly of vLAMPS.

For the standalone vLAMPS assembly, the three glass layers were secured in the aluminum clamp. The bottom side of the membrane forms the vascular chamber, and the top side forms the hepatic chamber (Supplementary Fig. **S1A**). The parallel perfusion of the 2 chambers was set up by two 10ml syringes in via a syringe pump (Harvard Apparatus, cat. no. PHD ULTRA) and connected via PEEK tubing (IDEX, cat. no. 1569XL) into the vLAMPS inlets. Efflux collection of ∼1.7ml was carried out in channel-independent (hepatic and vascular chamber) 4ml glass vials.

### PANIS fabrication

Prior to encapsulation into the PANIS, islets were washed twice in NF medium and equilibrated overnight at 37°C and 5% CO_2_. With the aid of a dissection microscope, ∼100 individual islets (per PANIS) were transferred using a 200 µL pipette (Fisher Scientific, cat. no. 02-707-430) into a suspension of 1.1% alginate solution. In 2 µL of the islet-alginate solution ∼10-20 islets were dispensed into the oval-well of the middle layer in an alternating array of ∼10 droplets. The alginate was polymerized by adding the solution of 10mM BaCl_2_ (Sigma, cat. no. 1.01716) through the bottom side of the membrane for 20 minutes at 37°C and 5% CO_2_, encapsulating the islets into each alginate bead. Lastly, the middle layer was fully submerged in NF (or EMS for disease studies) medium and placed at 37°C and 5% CO_2_ until assembly of PANIS.

Similar to the vLAMPS, the standalone PANIS assembly consisted of clamping the three glass layers in the aluminum clamp to form the top and bottom chambers (Supplementary Fig. **S1B**). The PANIS was placed under flow (35µl/hr) of medium from a single 10 mL syringe connected by PEEK tubing into the single PANIS inlet top chamber, and a loop was created by connecting the top chamber outlet to the bottom chamber inlet using C-flex tubing (Fisher Scientific, cat. no. NC0142679). The ∼1.7 mL efflux was collected from the bottom chamber outlet.

### Physical coupling

The 4-device aluminum clamp was used to form a duplicate coupling of a vLAMPS and a PANIS. Assembly of the vLAMPS and PANIS follows the same steps as described above, with coordination between vLAMPS cell seeding and PANIS islet incubation (Supplementary Fig. **S1D**). Two vLAMPS modules in parallel to each other were assembled first, followed by the assembly of the two PANIS modules. The vLAMPS was connected to the PANIS via C-flex tubing from the hepatic chamber (vLAMPS outlet number 8) into the PANIS top chamber (inlet number 5) (Supplementary Fig. **S1C**). With the system sealed, media perfusion was initiated in the inlets of the vLAMPS (inlet number 2 and 5) by connecting two 10 ml syringes to the device with PEEK tubing. Perfusion passed through the vLAMPS to the PANIS and followed the looped path as described above. Finally, the bottom chamber of the PANIS (outlet number 2) was fed into a collection tube.

### Efflux collection

Efflux medium from the vLAMPS, PANIS, and coupled vLAMPS-PANIS was collected in glass vials which were changed every other day for 8 days after initiating flow. Samples for the standalone vLAMPS were collected from the hepatic chamber (**Fig. S1A**), standalone PANIS samples were collected from the bottom chamber (**Fig. S1B**), and lastly medium collected for the coupled system was from the shared flow between the vLAMPS’s hepatic chamber, into PANIS top chamber, and looped to the bottom chamber (**Fig. 1C**). The collected samples were maintained at 4°C until the completion of the experiment. Efflux samples were then aliquoted in 500 µl samples and stored at −20 °C until analysis.

### Live and immunofluorescence islet imaging

Islet viability and morphology were evaluated after 8 days under perfusion. The islet-alginates beads were divided into four groups, one for live imaging, one for oxidative stress, one for immunohistochemistry, and one for glucose stimulated insulin secretion (GSIS) evaluation. The islets were decapsulated from the alginate by adding 100 mM EDTA and incubating for 2 minutes at room temperature. Viability was assessed by live/dead assay (Invitrogen, cat. no. L3224) following the manufacturer’s instructions. Imaging was done using Olympus widefield fluorescence microscope with 494/517nm and 528/617nm (excitation/emission) filters at 20x magnification. Viability in islets was determined by calculating the ratio between fluorescence intensity of live-cell area to the sum fluorescence intensity of live and dead cell area with both normalized to total number of nuclei, using Fiji (ImageJ version 1.53c).

The second subset of islets was evaluated for oxidative stress by 2’,7’-Dichlorodihydrofluorescein diacetate (DCFDA) (Abcam, cat. no. ab113851) staining of reactive oxygen species (ROS). Freshly decapsulated islets were washed with 1x buffer supplied in the kit and later transferred to a 20 μM DCFDA solution and incubated at room temperature for 1 h before a final wash with 1X buffer. To capture the total dimensions of each islet, we counterstained the islets with 1 μM MitotrackerTM Red CMX ROS (Invitrogen, cat. no. ab112143). Imaging was carried out on an Olympus Fluoview laser scanning confocal microscope (Olympus, Japan) with 490/520nm and 485/535nm (excitation/emission) for Mitotracker and DCFDA, respectively (20x magnification). For the ROS positive control, islets were preincubated with 50 μM of Tert-Butyl hydroperoxide (TBHP) (Sigma, cat. no. 416665) for 1 h before staining with DCFDA and Mitotracker. Image processing and quantification was carried out using Fiji. ROS in the islets was measured by quantifying the intensity of DCFDA signal in islets normalized to the area^89^. To account for islet size variability and the associated inconsistent signal to noise ratio of the islet’s core, the fluorescence intensity measurement was limited to the *periphery* of each islet. The *periphery* area was delineated as the distal one-third of the radius of each islet. The mean intensity in the selected region was normalized to the established *periphery* area of each islet. Normalized ROS intensity for islet groups was represented as a fraction of the corresponding value for the islets treated with TBHP (positive control). Area and intensity were measured using Fiji.

The third subset of released islets was fixed with 4% Paraformaldehyde (PFA) (Thermo Scientific, cat. no. J19943.K2) for 1 hour at room temperature. Following the previously descried methodology^47^, islets were immunofluorescence labeled using primary C-peptide (R&D Systems, cat. no. MAB80561) and glucagon (Abcam, cat. no. ab92517) antibodies with a 1:100 dilution. The primary antibodies were incubated at 4°C overnight. The islets were subsequently incubated at 4°C overnight with anti-mouse Alexa Fluor 488 (Invitrogen, cat. no. A28175) and anti-rabbit Alexa Fluor 555 (Invitrogen, cat. no. A27039) at a dilution of 1:500, and nuclei were stained using DAPI (1:2000) (Thermo Fisher, cat. no. D1306). Before imaging, islets were whole mounted on depression glass slides using Gold Antifade Mountant (Invitrogen, cat. no. P36930) and imaged using the Fluoview 1000 laser scanning confocal microscope with 40x objective with 1.32 NA. The images were acquired with an exposure time of 10 μs/pixel, and a line Kalman filter was applied to remove background noise. The images acquired for C-peptide, glucagon were quantified using a basic machine learning algorithm executed in Qupath (version 0.5.0) to detect the number of cells expressing C-peptide and glucagon.

### Glucose-stimulated insulin secretion

The fourth set of islet-alginate capsules was not decapsulated. Glucose stimulated insulin secretion (GSIS) was conducted under static conditions using 8 μm-membrane transwells (Corning, cat. no. 3464). Islet-alginate capsules were washed five times with G55 medium (500 mL HAMS F10 (Cytiva, cat. no. SH30025.01), 500 mL DMEM without glucose (Gibco, cat. no. A1443001), 600 mg NaHCO3, 110 mg CaCl2, and balanced to pH 7.4) prior to incubating overnight at 37°C and 5% CO_2_. Following the incubation, the islets were washed five more times and exposed to the low (3mM) and high glucose (16mM) G55 solutions for 1-hour spans. Insulin was measured from supernatant using an ELISA kit (ALPCO, cat. no. 80-INSHU-E01.1) as per the manufacturer’s instructions.

### Secretome measurements

Albumin, urea, LDH, COL1A1, and TIMP1 were measured as previously described^36,90^. Albumin assays were performed in a 1:100 efflux dilutions by enzyme linked immunosorbent assay (ELISA) using commercial antibodies (Bethyl Laboratories, cat. no. A80-129A and A80-129P) and an ELISA accessory kit (Bethyl Laboratories, E101) with a human albumin standard prepared in house (MilliporeSigma, cat. no. 126658). COL1A1 was measured using the Human pro-collagen 1A1 ELISA kit (R&D Systems, cat. no. DY6220-05) in a 1:50 efflux dilution, and TIMP1 was measured using the TIMP1 ELISA kit (R&D Systems, cat. no. DTM100) with a efflux dilution of 1:20. Urea was measured using the Stanbio BUN liquid reagent for diagnostic set (Stanbio Laboratory, cat. no. SB-0580-250). Lactate dehydrogenase was measured using the CytoTox 96 Non-Radioactive Cytotoxicity Assay (Promega, cat. no. G1780). The protocols for urea and LDH assay were modified to a 384 well microplate format with no efflux dilution. Controls were run in duplicate, and only sample values inside the range of the standard curve were used for analysis.

### Multiplex immunoassay

Amylin, C-peptide, Pancreatic Polypeptide, GLP-1, and glucagon were measured using a customized MILLIPLEX human metabolic Hormone Panel V3 (Millipore Sigma, cat. no. HMH3-34K). CCL2, CCL3, CXCL1, Fetuin A, G-CSF, GM-CSF, HGF, IGFBP-7, IL-1β, IL-6, IL-8, PDGF-AA, RBP4, TNF-α, and VEGF-A were measured using a custom Luminex Human Discovery LXSAHM-15 panel (R&D Systems). The measurements were carried out according to the manufacturer’s instructions at the University of Pittsburgh Cancer Proteomics Facility Luminex® Core Laboratory using the xMAP platform. All samples and controls were run in triplicate and only values inside the range of the standard curve were used for analysis.

### vLAMPS immunofluorescence imaging

On day 8 the vLAMPS cells were fixed with 2% PFA at 4°C for 30 minutes. Following the previously described methodology^36^, hepatocytes were immunofluorescence labeled for either cytokeratin 8 (CK-8) (1:200) (Invitrogen, cat. no. MA1-06318), or insulin receptor substrate 2 (IRS-2) (1:250) (Abcam, cat. no. ab134101). Stellate cells were labeled for alpha-smooth muscle actin (α-SMA) (1:100) (Sigma, cat. no. A2547). The primary antibodies were incubated at 4°C overnight. Secondary antibodies anti-mouse Alexa Fluor 488 (Invitrogen, cat. no. A-11001) and anti-rabbit Alexa Fluor 594 (Invitrogen, cat. no. A-11012) were incubated at room temperature for 2 hours (1:250). Lipid droplets in the hepatocytes were stained with LipidTOX (1:500) (Invitrogen, cat. no. H34476) fluorescent staining (595/615nm). The F-actin of LSECs were fluorescence stained with 568-Phalloidin (1:500) (Invitrogen, cat. no. A12380). Cell nuclei were stained using Hoechst (1:2,000) (Thermo Fisher, cat. no. 62249).

To capture the appropriate metabolic zonation across the vLAMPS, images were taken at the specific fields in the ROI denoted *Zone 1*, *Zone 2*, and *Zone 3* across the vLMAPS oval area (**Fig. S2E**). Image stacks were taken in a grid of 2 by 4 fields (length x height for a total of 8 fields per ROI). Images were acquired with a Nikon 20x (0.45 NA) objective using the Operetta CLS High Content Scanning system (Perkin Elmer). Applying confocal mode and maximum projection acquisition mode, all images were taken with at 3.3µm stack size on a 160µm height, using the 490/525nm, 561/572, and 650/671nm excitation/emission settings. Fluorescence intensity analysis was performed using Fiji. To exclude background fluorescence for IRS-2 and α-SMA the images were segmented with an intensity threshold based on a secondary antibody only control. To measure the total fluorescence intensity of the lipid droplets, particle quantification was applied by excluding small objects (<10 μm^2^). The average fluorescence intensity of all fields (regardless of the metabolic zone) was used for statistical normalization and analysis of LipidTOX, IRS-2, and α-SMA.

### Reproducibility analysis

Data from all studies were uploaded to the Numa Biosciences EveAnalytics™ platform. Inter-study reproducibility analysis followed the Pittsburgh Reproducibility Protocol (PReP)^56^ and used the Graphing/Reproducibility feature in the EveAnalyticsTM platform for statistical comparisons. Reproducibility status based on one-way ANOVA ranged from *Excellent* (p-value≥0.1) to *Poor* (p-value <0.01)^54,55^. All metrics with a p-value ≥0.01 were used for the multivariate correlation analysis.

### Multivariate correlation analysis

The correlation between the levels of the various secreted factors in the vLAMPS-PANIS coupled model was assessed using the Multivariate method analysis in JMP (version 17.1). The input data were the levels of the secreted analytes measured in the effluent from the bottom chamber of vLAMPS-PANIS coupled model. The correlation of each target analyte to each of the other target analytes measured under the same media conditions (NF or EMS) was estimated by the Pairwise method to calculate the Pearson correlation coefficient and the correlation probability p-value. The p-Value is the probability that the true correlation between two analytes is zero. Correlations with p-values <0.05 were considered to be statistically different from zero, i.e., a true correlation.

To verify for vLAMPS-derived and islet-derived factors, the levels of detection of the corresponding analyte had to be at least of 2.5:1 to the same factor under their corresponding medium between standalone vLAMPS (Supplementary **Fig. S4**) and standalone PANIS (Supplementary **Fig. S6**).

For the selection criteria of the key analytes’ correlation presented in **Table 2** the following criteria were used: paired analytes with reproducible p-values (>0.01) in EMS and NF medium, for one or both of the correlation coefficient p-values to be significant (≥0.05), and the magnitude of the significant correlation coefficients to be at least moderately correlated (r>0.6).

### Statistical analysis

Statistical comparisons between NF and EMS media groups as well as standalone and coupled systems were made using PRISM (version 10.1.2). All data are presented as mean ± standard error (SEM) for n= 2–7 devices per study. Normality was first determined with all data by Shapiro-Wilk test. When at least one of the groups or studies did not follow Gaussian distribution, a nonparametric test was applied. For media comparison studies, EMS values were normalized to the average NF values of their corresponding study. For parametric measurements a One-sample t test was applied, and a Wilcoxon signed-rank test was applied as nonparametric alternative. The two-way ANOVA with Sidak’s test was applied to multiple day efflux comparison between NF and EMS. An unpaired two-tail t test with Welch’s correction was applied to islet data collected outside of the PANIS, as no equal distribution and sample size could be secure due to the heterogeneric nature of islets derived from 3 different donors (Supplementary **Table S3**). Alternatively, Mann-Whitney test was implemented as the nonparametric test for one-to-one comparison of two unpaired groups.

## Data and metadata availability

All of the data and metadata were uploaded in EveAnalytics (previously BioSystics-Analytics Platform and MPS-Database).

## Acknowledgement

This work was supported by the following grants from the National Institute of Health (NIH) and National Science Foundation (NSF): UH3TR003289-NIH/NCATS (J. Behari, A. Soto-Gutierrez, and D.L. Taylor), U24TR002632-NIH/NCATS (M.E. Schurdak, A. Gough, and D.L. Taylor), UH3DK119973-NIH/NIDDK (A. Soto-Gutierrez, I. Banerjee, and D.L. Taylor), U24TR002632-NIH/NCATS (M.E. Schurdak, A. Gough, and D.L. Taylor), S10OD028450- NIH/OD (D.L. Taylor), R01DK117881-NIH/NIDDK (D.L. Taylor), 1706674-NSF/CBET (I.Banerjee), and 2229156-NSF/CBET (I. Banerjee). The authors would like to acknowledge valuable discussions with members of the Drug Discovery Institute and the Pittsburgh Liver Research Center (PLRC). We would like to thank the Biostatistical Consulting Laboratory at the University of Pittsburgh School of Public Health for its assistance with correlation analysis; however, any errors of fact or interpretation remain the sole responsibility of the authors. The cytokine data presented here were obtained using the UPCI Cancer Biomarkers Facility: Luminex Core Laboratory.

## Author contributions

J.A., R.K., C.W., and L.V. conceptualized and designed the experiments under the supervision of A.G., V.Y., M.T.M., A.M.S., I.B., and D.L.T. J.A., R.K., and C.W. participated in MPS experiments. J.A., R.K., C.W., D.G., C.R., R.D., and G.L. participated in the efflux and imaging quantification. J.A., M.E.S., and D.A. participated in reproducibility and correlation analysis. A.S.G., J.B., and V.J. lead the clinical input. J.A., R.K., C.W., M.E.S., M.T.M., and A.M.S. wrote the manuscript under the supervision of V.Y., I.B., and D.L.T. All authors have read and agreed to the published version of the manuscript.

## Corresponding authors

Correspondence to Andrew M. Stern, Ipsita Banerjee, and D. Lansing Taylor

## Abbreviations

α-SMA: alpha-actin, alpha-actin-2
CCL2: C-C motif chemokine ligand 2
CCL3: C-C motif chemokine ligand 3
COL1A1: Collagen type I alpha 1 chain
CXCL1: C-X-C motif chemokine ligand 1
EMS: Early metabolic syndrome
G-CSF: Granulocyte colony-stimulating factor
GLP-1: Glucagon like peptide 1
GM-CSF: Granulocyte-macrophage colony-stimulating factor
GSIS: Glucose-stimulated insulin secretion
HGF: Hepatocyte growth factor
IGFBP-7: Insulin like growth factor binding protein 7
IL-1β: Interleukin 1 beta
IL-6: Interleukin 6
IL-8: Interleukin 8
ISI: Insulin stimulation index
NF: Normal fasting
IRS-2: Insulin receptor substrate 2
PDGF-AA: Platelet-derived growth factor subunit A
PP: Pancreatic polypeptide
RBP4: Retinol binding protein 4
ROS: Reactive oxygen species
TIMP1: TIMP metallopeptidase inhibitor 1
TNF-α: Tumor necrosis factor alpha
VEGF: Vascular endothelial growth factor A

## Supplementary figures

**Supplementary Table. S1.** Media composition for the normal fasting and early metabolic syndrome states used in both standalone and coupled MPS.

| Component | NF<br>(Normal Fasting) | EMS<br>(Early Metabolic Syndrome) |
| --- | --- | --- |
| Basal | William's E w/o glucose | William's E w/o glucose |
| HEPES | 15 mM | 15 mM |
| Dexamethasone | 100 nM | 100 nM |
| Pen/Strep | 0.5% | 0.5% |
| FBS | 1% | 1% |
| Gluta-MAX | 1x | 1x |
| BSA | 1.25 mg/ml | 1.25 mg/ml |
| Linoleic acid | 5.35 µg/ml | 5.35 µg/ml |
| Transferrin | 6.25 µg/ml | 6.25 µg/ml |
| Sodium selenite | 6.25 ng/ml | 6.25 ng/ml |
| Glucose | <u>5.5 mM</u> | <u>20 mM</u> |
| Insulin | <u>10 pM</u> | <u>10,000 pM</u> |
| Glucagon | <u>100 pM</u> | <u>10 pM</u> |
| Oleic acid | - | <u>200 µM</u> |
| Palmitic acid | - | <u>100 µM</u> |

Media modulation of key metabolites (glucose, insulin, glucagon, FFAs) allows replication of progressive MASLD. Media formulations were designed based on physiologically relevant metabolites levels in plasma during normal fasting (NF) and early metabolic syndrome (EMS)<sup>58</sup>. Underlined terms indicate differences in key media components in the NF and EMS media formulations.

**Supplementary Fig. S1.**
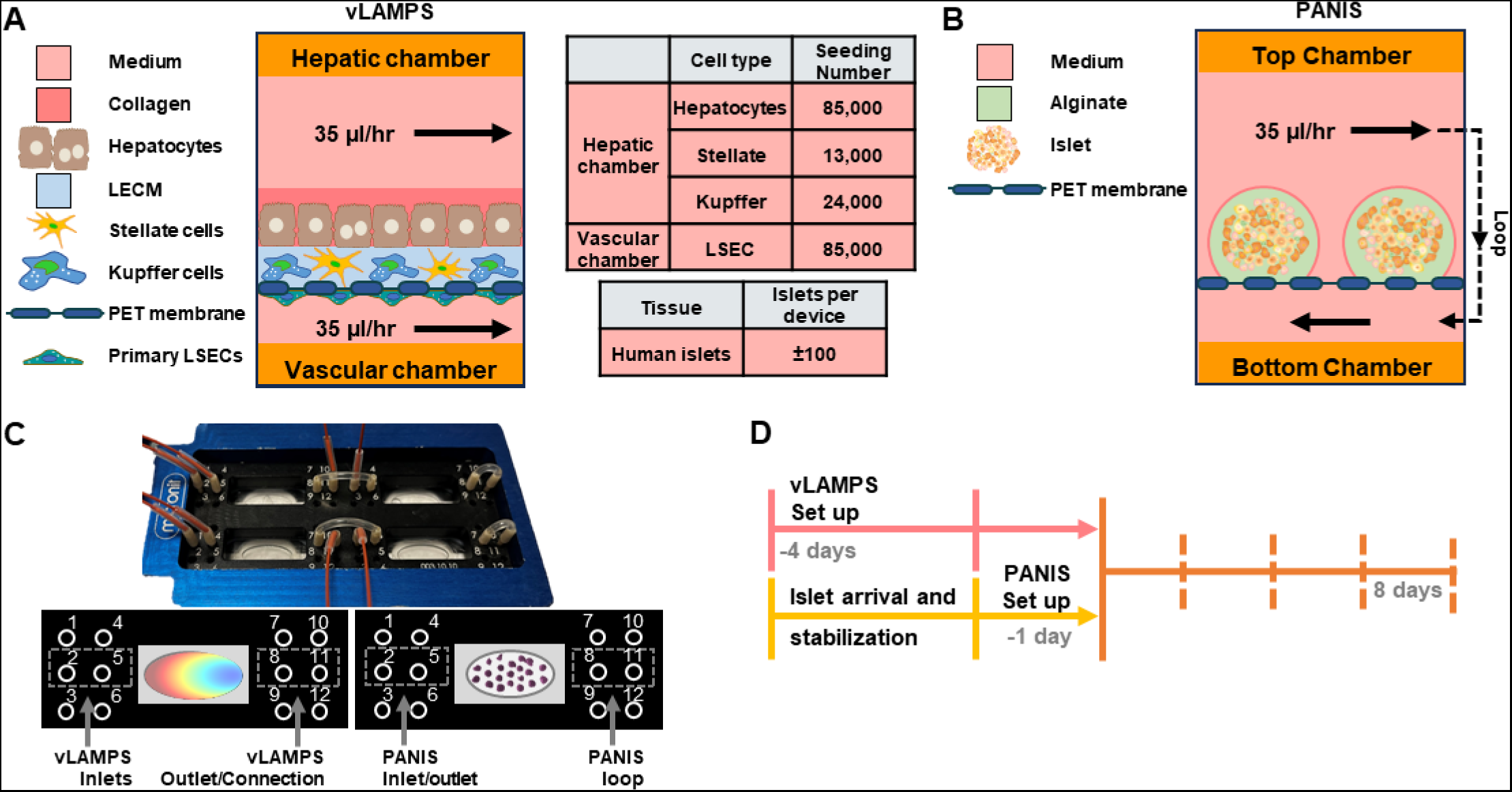
Standalone vLAMPS and PANIS are designed to recapitulate critical physiological aspects of the liver and pancreas and enable the preservation of their complex physiology during coupling. (**A**) Diagram of the vLAMPS showing the biomimetic cell composition and arrangement, the two compartments (hepatic and vascular), and the direction of the parallel flow at 35µl/hr. (**B**) Diagram of the PANIS, and the flow path that initiates in the top chamber and continuing through a loop into the bottom chamber. The islets are encapsulated in an array of alginate-based capsules. The shared flow rate for both systems was selected to first achieve the liver’s metabolic oxygen zonation (**Fig. S2**), and second maintain islets viability (**Fig. S5**). (**C**) Representative picture of the coupled system and fluidic arrangement between vLAMPS and PANIS in the four-chamber system. (**D**) Timeline for setup of each standalone system prior to coupling, followed by 8-day single pass perfusion with medium collection every other day.

**Supplementary Fig. S2.**
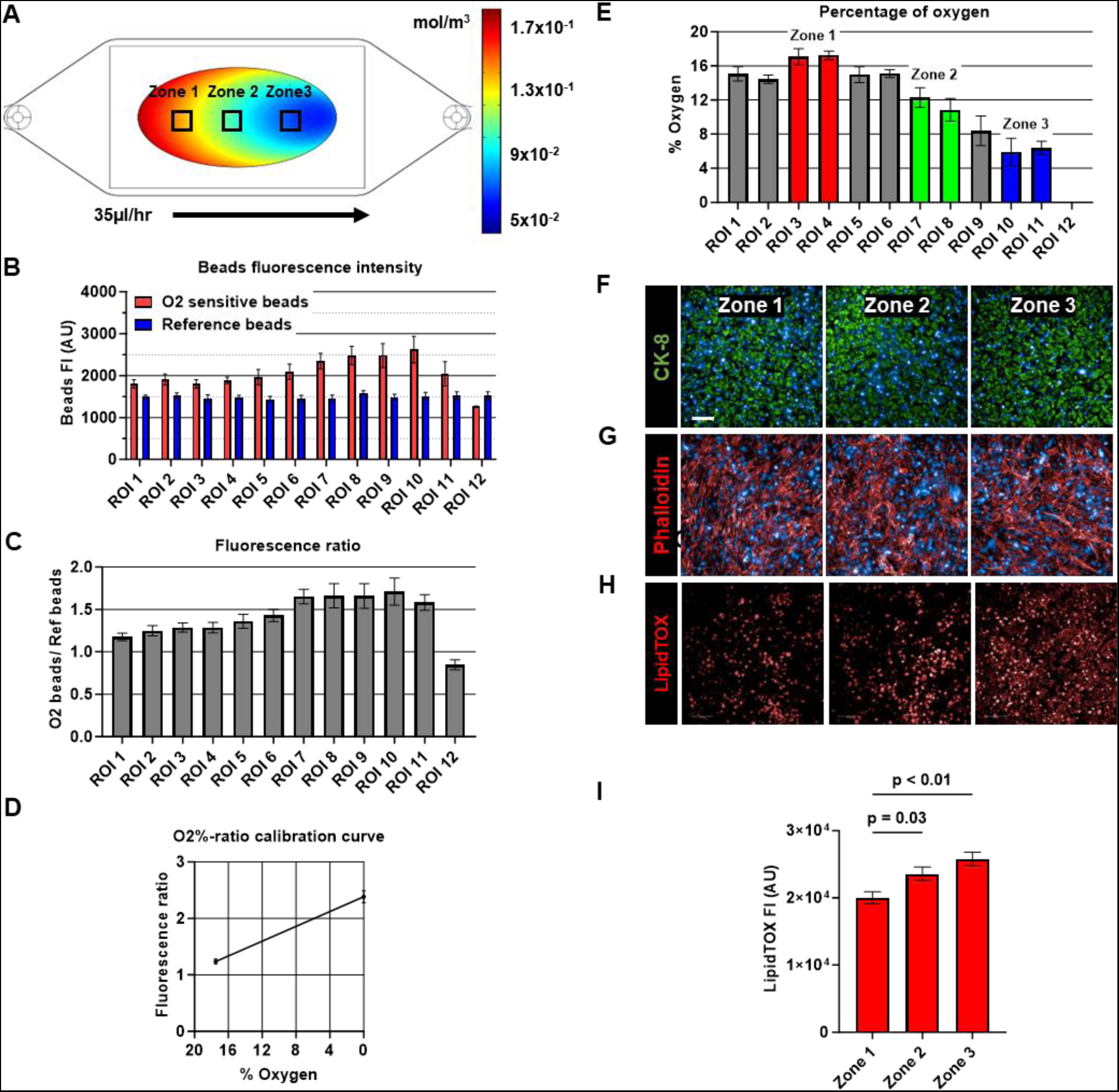
The vLAMPS exhibits metabolic oxygen zonation. The regions of interest (ROIs) were specific microscopic fields selected across the cell culture area, using the oxygen simulation as reference of candidate oxygen tension locations. (**A**) COMSOL Multiphysics-based model of oxygen tension with a concentration across the tissue substrate between 1.8×10^-1^ to 4×10^-2^ mol/m3. Representative liver acinus metabolic zones have been identified; zone 1, zone 2, and zone 3 are proximal, medial, and distal from the inlet of the two-chamber platform, respectively. (**B**) Fluorescence intensity (FI) of the oxygen sensitive and reference beads placed in the hepatic chamber of the vLAMPS. The measurement of the oxygen tension was carried out by ratiometric method^36^ by dividing the FI of the oxygen sensitive beads by the FI of the reference beads. (**C**) Fluorescence ratio of each ROIs. (**D**) The internally generated standard curve (see Methods) was used to determine the oxygen tension concentrations values from the ratios. (**E**) Percentage of oxygen tension measured across the ROIs, *zone 1* with an average oxygen tension of 17%±0.5 (ROIs 3-4, red), *zone 2* 11.5%±0.8 (ROIs 7-8, green), and *zone 3* 6%±0.8 (ROIs 10-11, blue). To determine the overall integrity of the cells within the chip under flow at the determined oxygen tensions, images at the ROIs corresponding to specific oxygen tensions were taken under NF medium conditions on day 8. Representative immunofluorescence images of labeled (**F**) hepatocytes (CK-8), (**G**) liver sinusoidal endothelial cells (Phalloidin), and (**H**) lipid droplets stained with LipidTOX in vLAMPS maintained in NF medium. Scale=100µm. (**I**) FI quantification of LipidTOX stained lipid droplets following 8-day perfusion. All bar graphs display the mean and standard error (SEM). The statistical analysis in graph - *I -* was done by one-way ANOVA and Bonferroni multiple comparison test to the mean of zone 1 with p-values shown.

**Supplementary Fig. S3.**
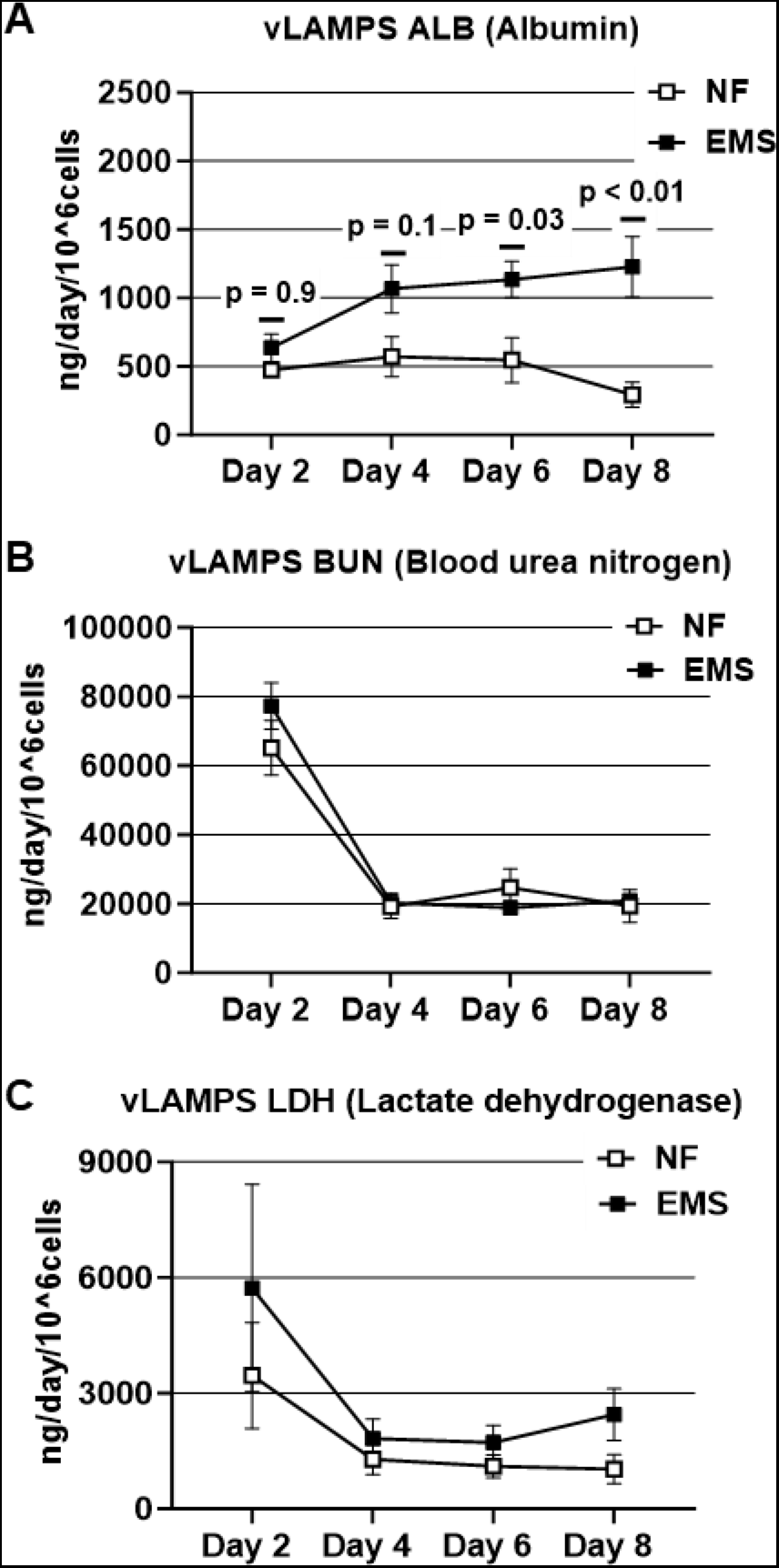
Basic liver functions were maintained in standalone vLAMPS under NF and EMS conditions during an 8 day period (supports Fig. 2). (**A**) Efflux from the hepatic chamber had approximately the same initial albumin concentration between both medias and demonstrated an increasing tendency in EMS medium over the experimental time course. (**B**) Urea secretions and (**C**) LDH secretions were consistent across the 8 days between NF and EMS. Line graphs display the mean and standard error (SEM). For graph *-A-* Statistical analysis was done by two-way ANOVA and Sidak’s multiple comparison test to the mean albumin secretion between NF and EMS with p-values shown.

**Supplementary Fig. S4.**
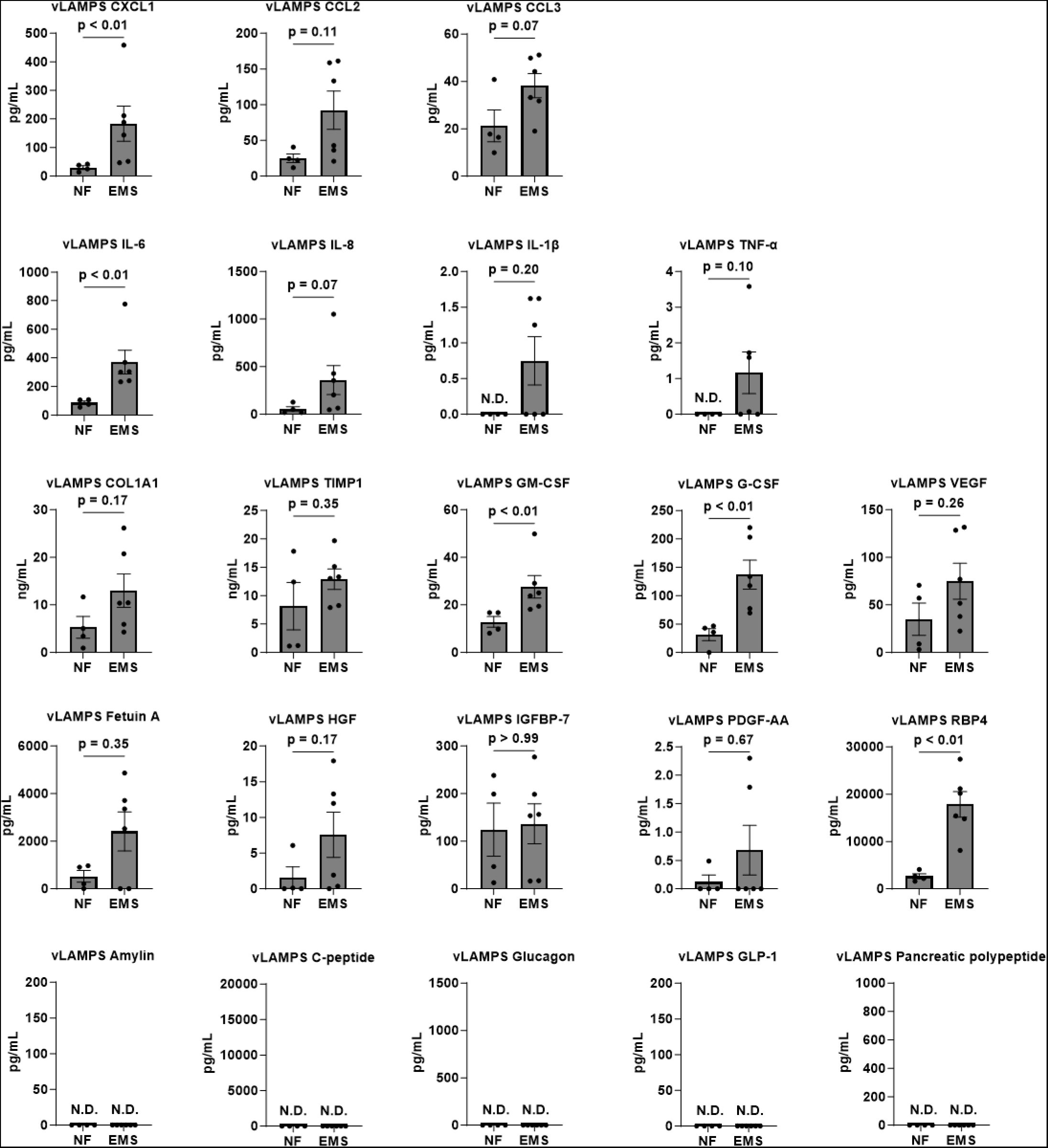
In comparison to NF conditions, EMS conditions perturb the standalone vLAMPS secretome at day 8 (supports Fig. 2 and Table 1). Chemokines, cytokines, fibrotic, pro-inflammatory, and hepatokines have elevated tendencies in EMS compared to NF. There was no detection of the islet derived factors Amylin, C-peptide, glucagon, GLP-1, and PP. Scatter plots represent the mean of all studies and SEM for NF and EMS. Statistical analysis between the media conditions were carried out by two-tail Mann-Whitney test with p-values shown. Bars with -N.D.- denotes *no detection* by assay because values were below the lowest value of the standard curve.

**Supplementary Fig. S5.**
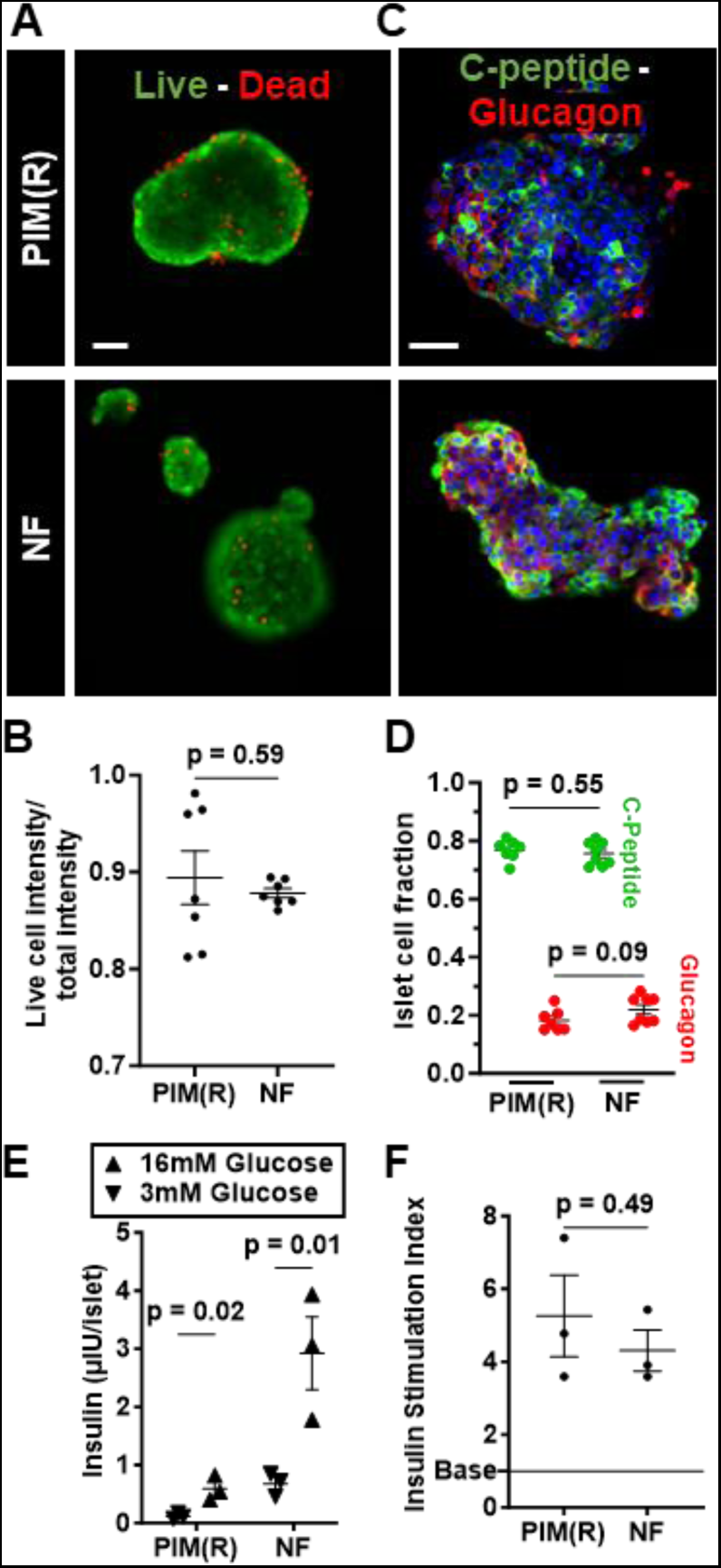
NF media supports the healthy phenotype and functionality of PANIS over 8 days of culture. Standalone PANIS were evaluated with NF medium as an alternative medium to the islet-exclusive PIM(R) medium. (**A**) Representative images of islets in the PANIS system for viability (live/dead staining) after culture in PIM(R) and NF media, respectively. (**B**) Viability quantification shows no significant difference between PIM(R) and NF. (**C**) Representative immunofluorescence labeling of C-peptide and glucagon in PIM(R) and NF media, respectively. All scale bar=50µm. (**D**) The fraction of quantified C-peptide and glucagon positive cells shows no significant difference between the two media. (**E**) GSIS response of islets cultured in PIM(R) and NF media and their (**F**) insulin stimulation indices showing no significant difference between PIM(R) and NF media. Scatter plot graphs display the mean and standard error (SEM) with p-values shown. For all graph statistical analysis was done by two-tail t test with Welch’s correction.

**Supplementary Fig. S6.**
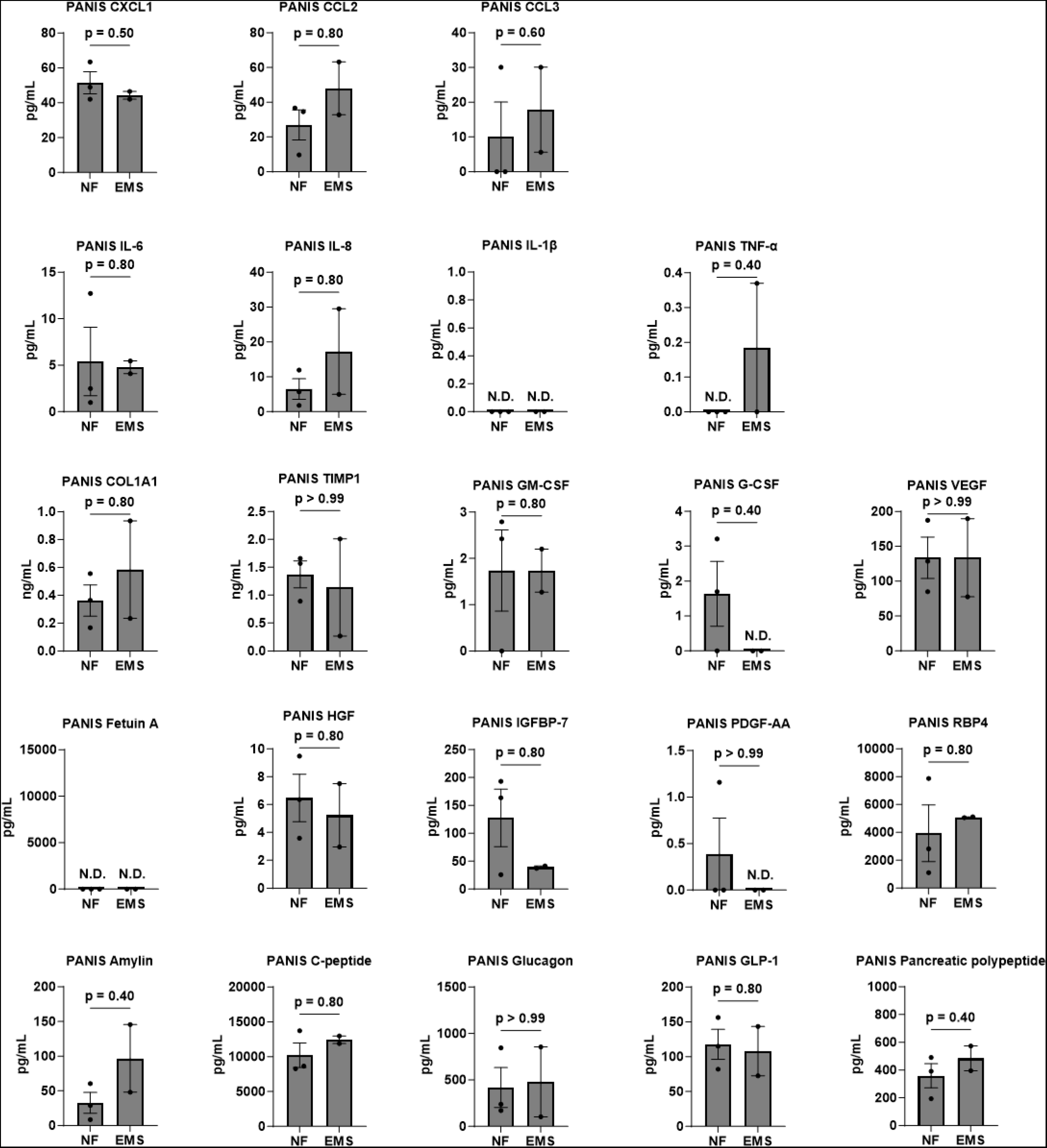
No significant differences in the level of secreted factors from the standalone PANIS perfused with NF or EMS media were observed at day 8 (supports. **Fig. 3 and Table 1).** The secretion of chemokines, cytokines, fibrotic, pro-inflammatory, and islet factors demonstrate no significant differences between NF and EMS. In the PANIS, efflux of some hepatokines were detected at noteworthy intensities including the levels of detection of CCL2, CCL3, IL-1β, VEGF, HGF, and PDGF-AA, are in the same range as the vLAMPS under NF and EMS. Fetuin A was the only analyte not detected in the PANIS that was detected in vLAMPS (Supplementary **Fig. S4**). Scatter plots represent the mean of all studies and SEM for NF and EMS. Statistical analysis between the media conditions were carried out by two-tail Mann-Whitney test with p-values shown. Bars with -N.D.- denotes *no detection* by assay.

**Supplementary Fig. S7.**
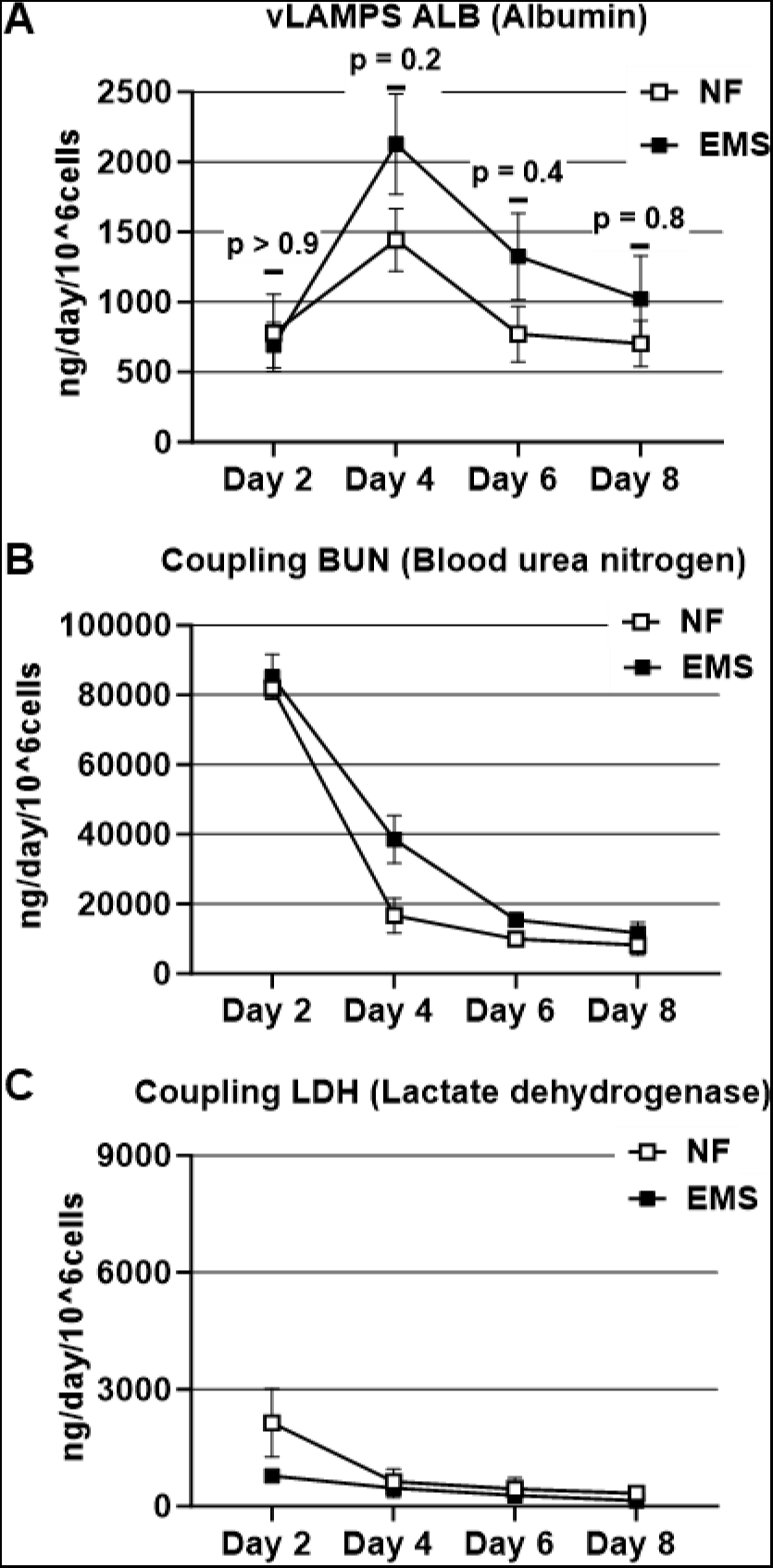
Basic liver functions were maintained in the coupled vLAMPS-PANIS, under both NF and EMS conditions. (supports. **Fig. 4).** (**A**) Efflux from the coupling chamber had approximately the same initial albumin concentration as the standalone vLAMPS (**Fig. S3A**) as well demonstrating a similar increasing trend under EMS. (**B**) Urea and (**C**) LDH secretions were also consistent with standalone vLAMPS (**Fig. S3B** and **S3C**). Line graphs display the mean and standard error (SEM). For graph *-A-* Statistical analysis was done by two-way ANOVA and Sidak’s multiple comparison test to the mean albumin secretion between NF and EMS with p-values shown.

**Supplementary Fig. S8.**
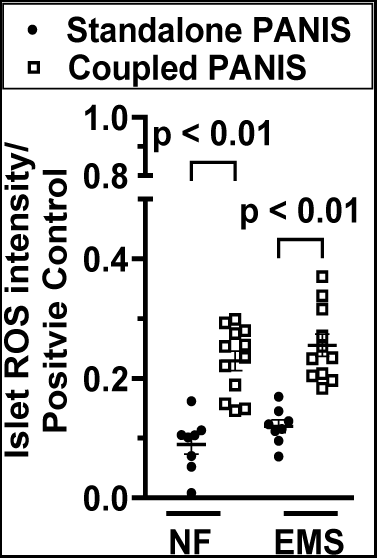
Independent of media conditions, islets have elevated ROS levels in the coupled compared to standalone modules. Comparison between ROS values of islets from standalone PANIS versus islets in the coupled vLAMPS-PANIS, with their respective media (NF or EMS). Scatter plot graph display the mean and standard error (SEM) with p-values shown. Statistical analysis by unpaired two-tail t test with Welch’s correction with p-values shown.

**Supplementary Fig. S9.**
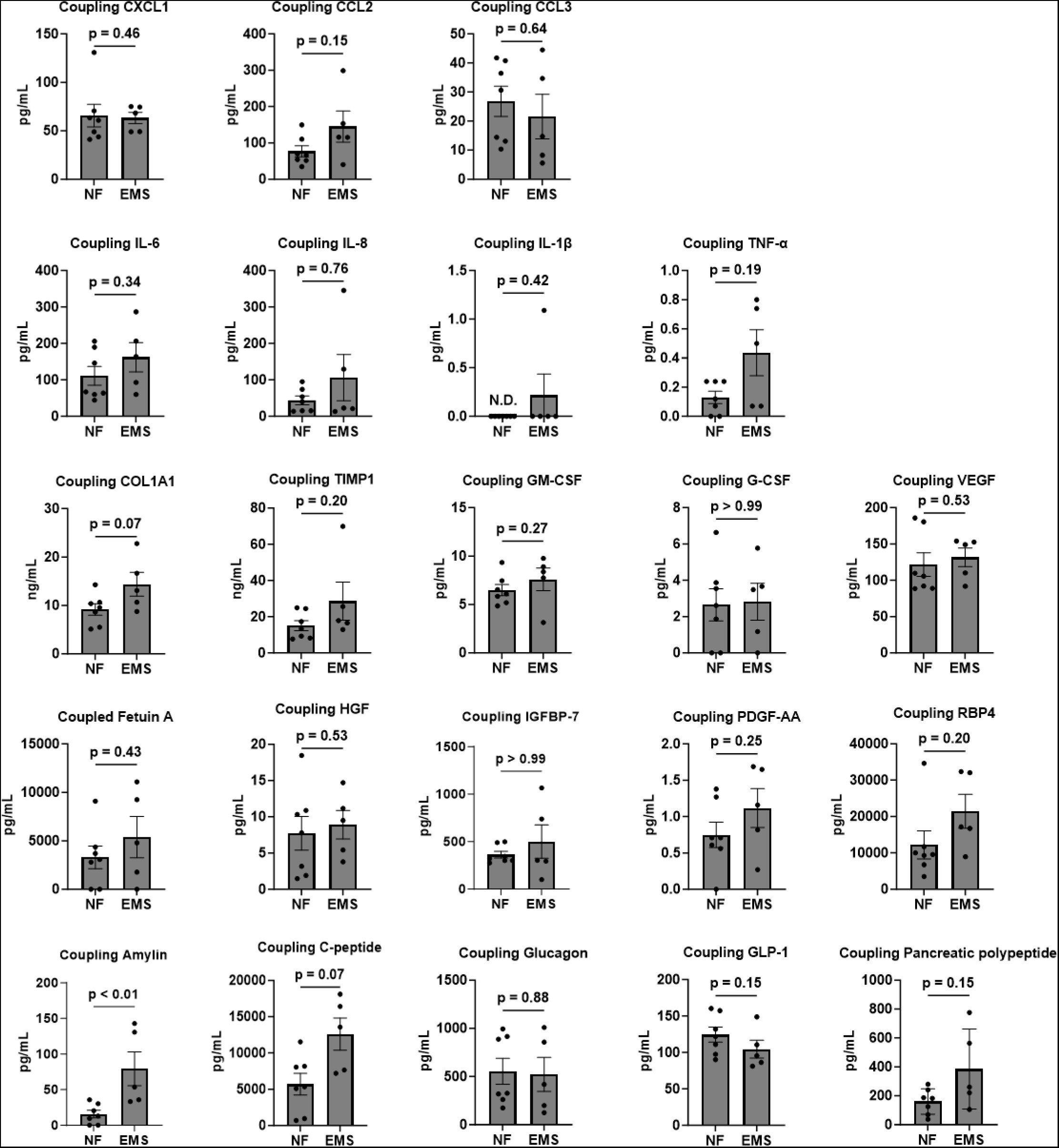
The level of islet derived factors in the coupled vLAMPS-PANIS efflux at day 8 are perturbed by EMS media relative to NF conditions. (supports Fig. 4). The secretion of islet-derived factors remained in the same range of detection as in the standalone PANIS; however, an increasing overall trend was observed in secretion of islet derived factors between NF and EMS, except for glucagon and GLP-1. The levels of detection of IL-6, COL1A1, TIMP1, Fetuin A, IGFBP-7, and RBP4, are in the same range as the standalone vLAMPS but above the range of detection of the standalone PANIS, in both NF and EMS conditions. Scatter plots represent the mean of all studies and SEM for NF and EMS. Statistical analysis between the media conditions were carried out by two-tail Mann-Whitney test with p-values shown. Bars with -N.D.- denotes *no detection* by assay.

**Supplementary Fig. S10.**
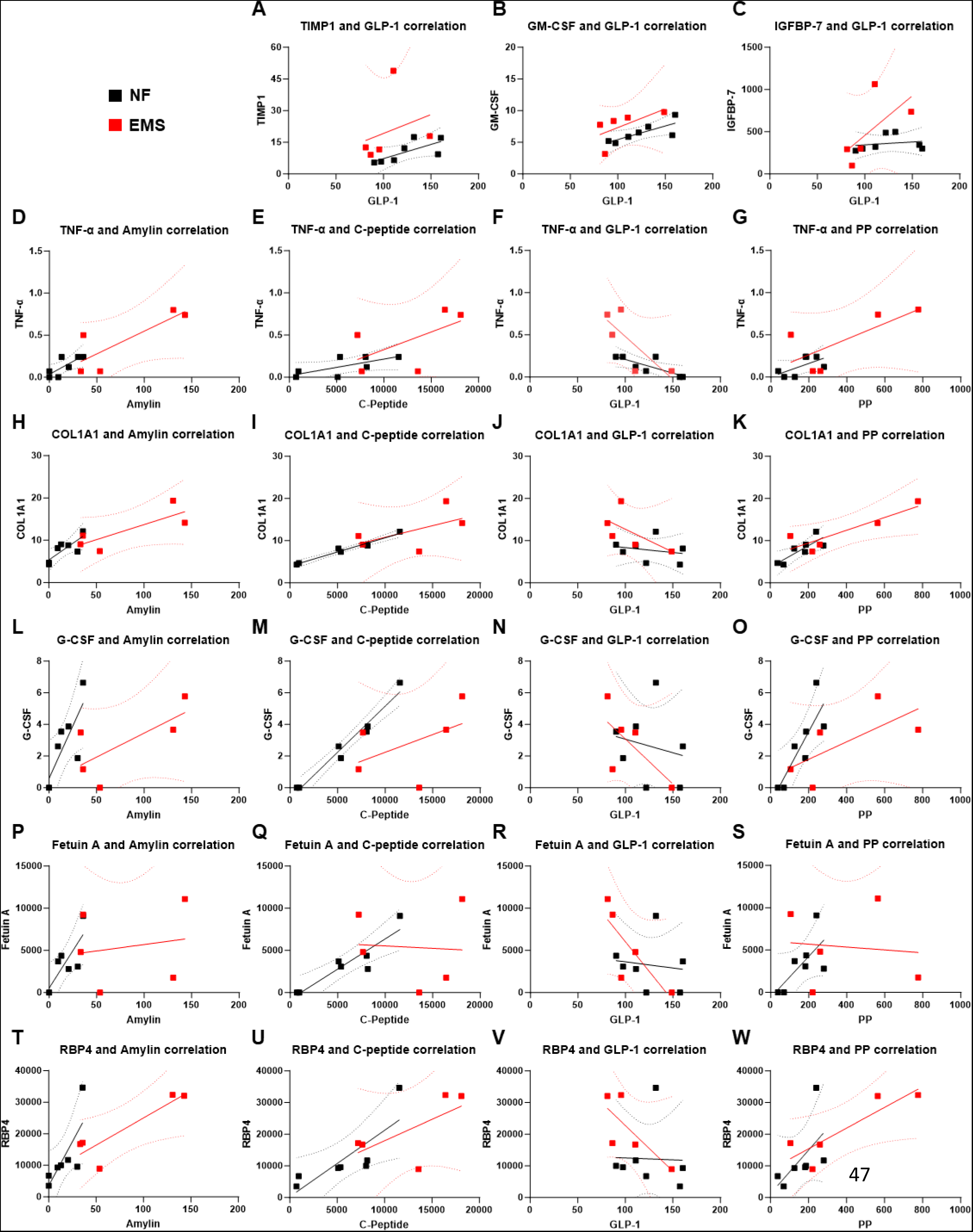
Pairwise correlations involving a vLAMPS derived factor and an islet derived factor in the coupled vLAMPS-PANIS efflux are altered under NF versus EMS conditions consistent with a role of hepatic factors in the dysregulation of islet function (supports Table 2). (**A**) TIMP1, (**B**) GM-CSF, and (**C**) IGFBP-7 have a significant reproducibility (p≥0.05) and positive correlations to GLP-1. TIMP1 and GM-CSF have a more significant correlation to GLP-1 under NF, but under EMS the correlation becomes significant for IGFBP-7 under EMS. TNF-α has a positive correlation to (**D**) Amylin under EMS, while for (**E**) C-peptide the opposite was observed. This contrasts with the significant negative correlation between (**F**) GLP-1 and the cytokine under NF. The same vLAMPS derived analyte has a positive correlation to the islet derived factor (**G**) pancreatic polypeptide in EMS. The liver fibrosis associated protein COL1A1 has a significant correlation positive to (**H**) Amylin and (**I**) C-peptide in NF. There is a reproducible (p≥0.05) and negative correlation with (**J**) GLP-1 in EMS. Equally reproducible, but significant positive correlation was detected with (**K**) pancreatic polypeptide in both media conditions. Similarly, G-CSF has a significant positive correlation to (**L**) Amylin and (**M**) C-peptide in NF. On the other hand, a negative correlation with (**N**) GLP-1 under EMS. With a significant positive correlation with (**O**) pancreatic polypeptide in normal fasting conditions. The vLAMPS exclusive Fetuin A has a strong and significant positive correlation to (**P**) Amylin and (**Q**) C-peptide, which completely gets lost in EMS conditions. Further, under EMS this hepatokine and (**R**) GLP-1 have a strong negative correlation. On the contrary, the correlation to **S** PP is a positive one in NF. The transporter protein RBP4 has a significant positive correlation with **T** Amylin in both media conditions. When correlated to (**U**) C-peptide the significant positive correlation is only observed in NF. A significant reproducibility (p≥0.05) and negative correlation to (**V**) GLP-1 was observed in disease conditions. Equally reproducible but with a significant positive correlation was the relationship between the vLAMPS-factor and (**W**) pancreatic polypeptide under the same EMS conditions.

**Supplementary Fig. S11.**
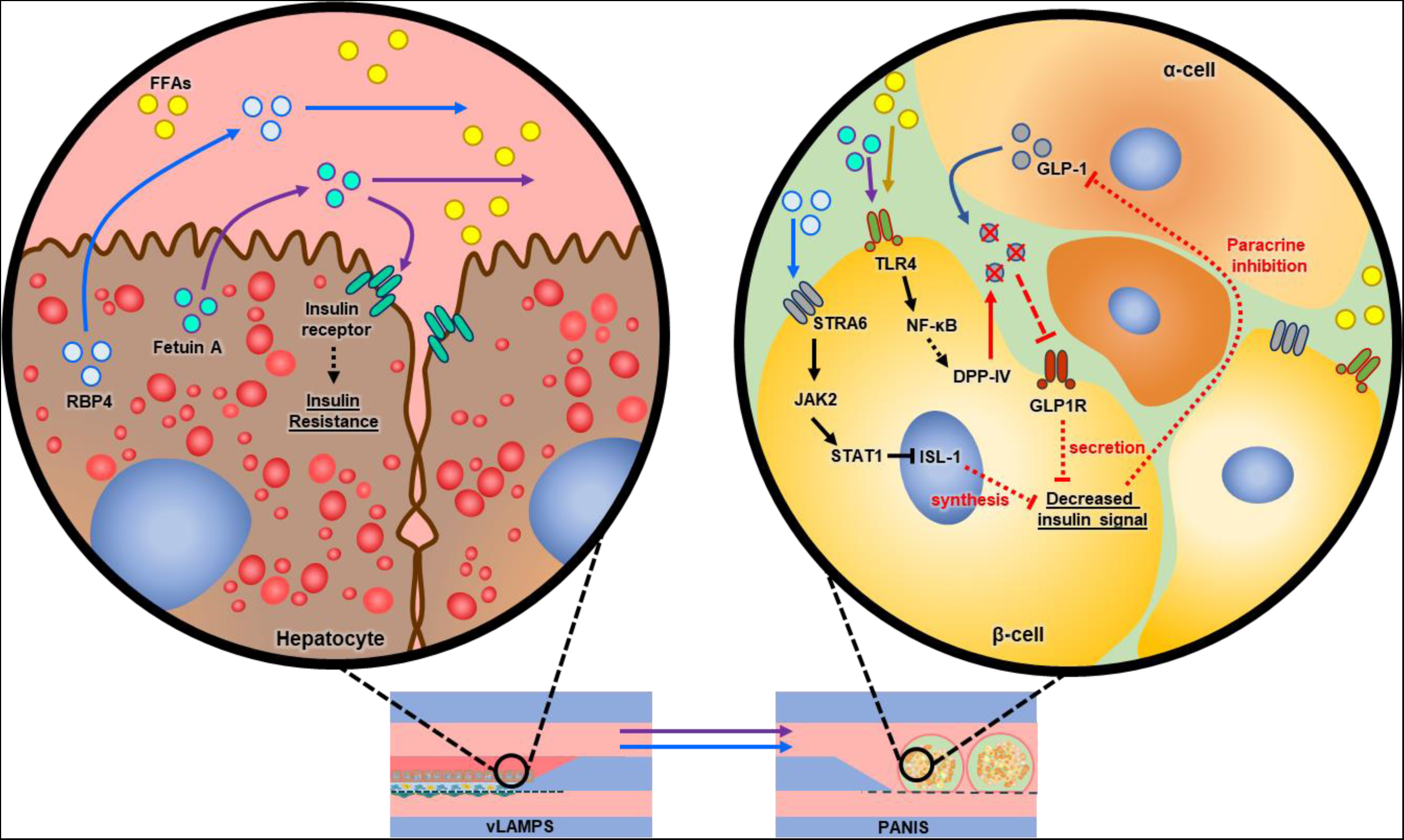
Schematic representation of the vLAMPS derived hepatokines RBP4 and Fetuin A negative impact on the pancreatic β-cell insulin response under EMS conditions. Hepatocytes with increased steatosis (red bubbles) have an elevated secretion of RBP4 (light blue beads) and Fetuin A (aqua beads) in the EMS medium with high FFAs (yellow beads) content. During fasting state (NF) Fetuin A contributes to glucose homeostasis by acting as a negative modulator of the insulin receptor. The steady stream of glucose is enough to stimulate insulin secretion during fasting state (positive correlation between Fetuin A/C-peptide under NF, supplementary **Fig. S10Q**). The higher levels of Fetuin A during EMS state may interfere with the hepatocyte’s insulin receptor activity potentiating the MASLD progression leading into insulin resistance. Downstream in the PANIS, these same Fetuin A levels alongside with FFAs promotes GLP-1 (grey beads) degradation through the TLR4-NF-κB-DPP-IV pathway. Truncation of GLP-1 by DPP-IV (negative correlation between Fetuin A/GLP-1 under EMS, supplementary **Fig. S10R**) decreases the insulin releasing actions of the GLR-1 receptor (GLP1R) in β-cells. Elevated RBP4 downregulates insulin synthesis in β-cells by the STRA6-JAK2-STAT1 pathway (decrease in of RBP4/C-peptide correlation under EMS, supplementary **Fig. S10U**). The decreased insulin signal could exacerbate the decreasing GLP-1 signal through paracrine inhibition. These processes lead to a decrease in the islet’s glucose sensitivity (decreasing GSIS) due to underlying impaired insulin response.

**Supplementary Table. S2.** Primary hepatocytes donor information.

| Thermo Fisher Lot # | Age | Sex | Ethnicity | BMI | CYP1A2 activity | CYP2B6 activity | CYP3A4 activity |
| --- | --- | --- | --- | --- | --- | --- | --- |
| HU1960 | 65 | f | African American | 27.12 | 550 | 53.8 | 992 |

**Supplementary Table. S3.** Islet donor information.

| Prodo Labs # | Age | Sex | Ethnicity | BMI | HbA1c | Height (in) | Weight (lb) |
| --- | --- | --- | --- | --- | --- | --- | --- |
| HP-23046-01 | 60 | m | Hispanic | 32.9 | 5 | 69 | 224 |
| HP-23054-01 | 46 | m | Hispanic | 27.4 | 5.6 | 65 | 164 |
| HP-23083-01 | 47 | f | Hispanic | 30.6 | 5.5 | 59 | 152 |

**Supplementary Table. S4.**
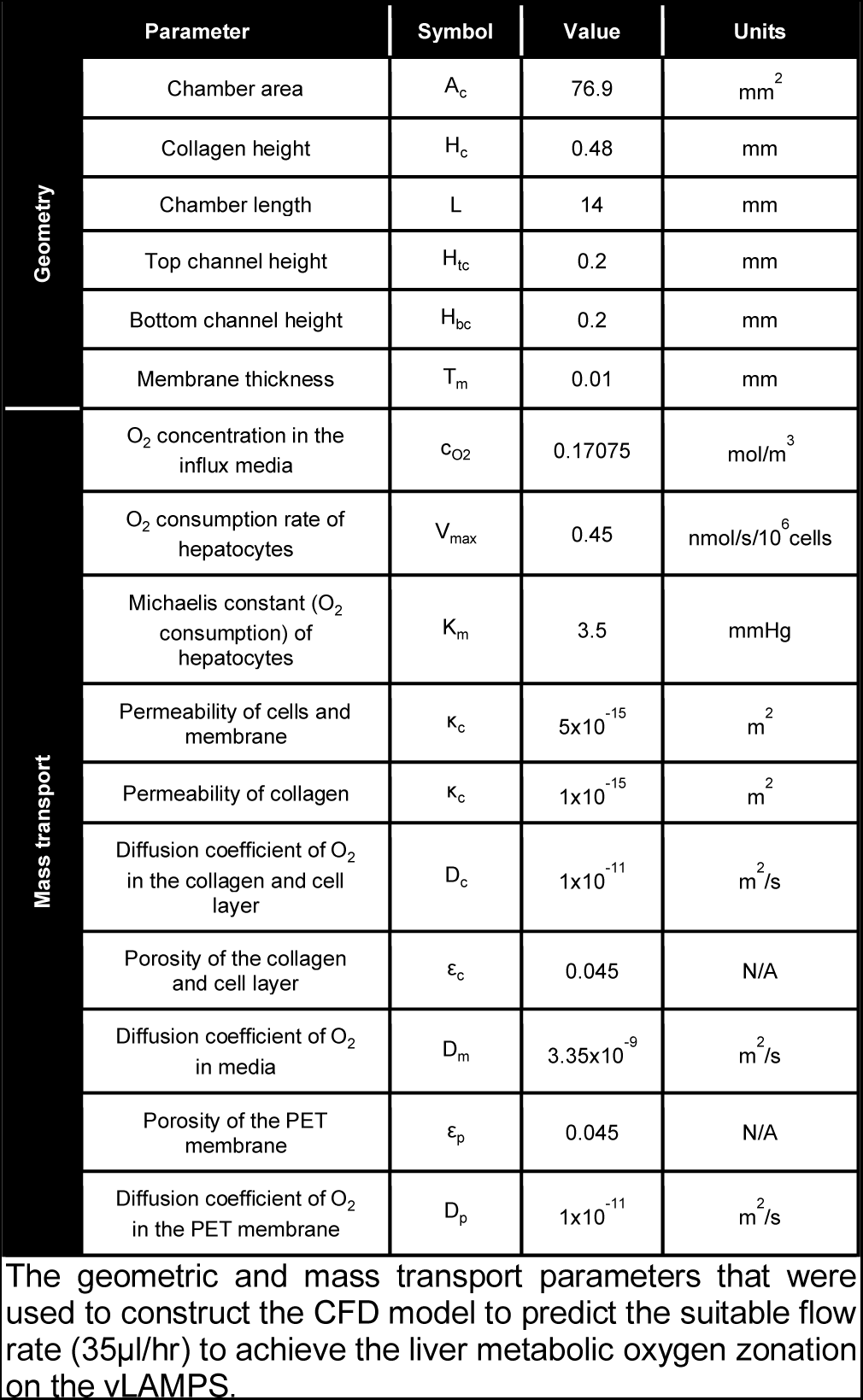
Parameters used in computational fluidic dynamics simulation to predict the O_2_ zonation in vLAMPS.

